# P53 is a direct regulator of the immune co-stimulatory molecule CD80

**DOI:** 10.1101/2021.05.24.445214

**Authors:** Eziwoma Alibo, Gurkan Mollaoglu, Maxime Dhainaut, Royce Zhao, Samuel Rose, Alessia Baccarini, Ramon Parsons, Brian D. Brown

## Abstract

Increasing evidence indicates oncogenes and tumor suppressors not only influence cell fitness but can also control the immunophenotype of cells. Here, we examined how 34 commonly mutated genes in colorectal cancer (CRC) may influence the expression of 8 key immunomodulatory proteins. To do this, we employed a functional genomics approach utilizing Pro-Code/CRISPR libraries for high-dimensional analysis. We introduced a library of 102 Pro-Code/gRNA combinations, targeting each of the 34 genes, in CT26 cells, a CRC cell model, and measured the expression of each of the immunomodulatory proteins by CyTOF mass cytometry. Notably, cells carrying a Pro-Code/CRISPR targeting the Trp53 lost expression of the immune co-stimulatory molecule CD80. Validation confirmed that Trp53 knockout resulted in the loss of CD80 and that activation of P53, through DNA damage or stabilization, resulted in CD80 upregulation. P53 ChIP-seq identified the CD80 promoter as a direct target of P53. CD80 regulation by P53 was identified in other cells, including normal epithelial cells and macrophages. Functionally, CD80 reduction caused by P53 loss led to a reduced capacity for CRC to prime antigen-specific T cells. These studies establish CD80, a canonical co-stimulatory molecule, as a direct target of the tumor suppressor and DNA damage response gene, P53.

## INTRODUCTION

Cancer cell interactions with T cells are highly dependent on the expression of several cell surface receptors and ligands. MHC class I (MHC-I), for example, is required for the T cell receptor (TCR) to engage cancer cells in an antigen-dependent manner^1^. Downregulation of MHC-I can enable cancer cells to escape T cell killing. While MHC-I engagement of the TCR is sufficient to activate memory T cells, naïve T cells also require the engagement of CD28 on their surface by co-stimulatory molecules, such as CD80 and CD86, on the target cell surface. In addition to molecules that are important for stimulating the immune system, cells can also upregulate molecules that subvert immunity and prevent their clearance. For example, cancer cells often upregulate checkpoint ligands, such as PD-L1 or CD47, which have immune-suppressive functions. PD-L1 acts by binding PD1 on activated T cells, and this induces T cell exhaustion. Whereas CD47 engages SIRPα on macrophages and serves as a ‘don’t eat me’ signal^2^. An important question in cancer immunology, and more broadly in immunology, is how each of the immunostimulatory and immunoinhibitory molecules, which make up the immunophenotype of a cell, is controlled.

Oncogenes and tumor suppressors regulate multiple facets of tumorigenesis, more notably cancer cell proliferation and survival. However, it has also become evident that classical cancer regulators (i.e. tumors suppressors and oncogenes) have a role in controlling the ability of tumors to evade immune clearance^3–5^. In some cases, cancer cell mutations modify the tumor immune microenvironment by decreasing the recruitment of anti-tumor immune cells or by increasing the recruitment of immunosuppressive cells. For example, activated WNT/β-catenin signaling decreases dendritic cell recruitment in melanoma^6^ and loss of NKX2-1 increases neutrophil recruitment in lung cancer^7^. In other cases, cancer cell mutations change the immunogenicity of the cancer cell more directly. Induction of the oncogene MYC, using a tet-ON system, resulted in upregulation of both PD-L1 and CD47^8^. Mutations in JAK and STAT render cells unresponsive to IFN-γ signaling which results in lower MHC-I levels and poor clinical response to drugs that inhibit the PD1/PDL1 pathway^9^.

As there are many oncogenes and tumor suppressors and different immunoregulatory molecules of relevance, identifying possible connections is not trivial. Pooled CRISPR screens have enabled proliferative and survival functions of cancer-associated genes to be identified at greater scale and speed than possible with candidate approach, but these screens are generally limited to identifying gene functions associated with cell fitness. This is because the readout of the screens is ‘enrichment’ or ‘de-enrichment’ of a particular CRISPR/knockout, and not the expression of specific molecules. To address this limitation, we recently developed a protein barcode (Pro-Code) technology that enables CRISPR screens to be read out using flow cytometry or CyTOF mass cytometry^10^. We created a library of Pro-Code expressing lentiviral vectors which each encodes a different CRISPR gRNA. When cells are transduced with the library, each cell expresses a unique Pro-Code and CRISPR. Using CyTOF, cells carrying a Pro-Code/CRISPR, and thus having a particular gene knockout, can be resolved. In addition to detecting the Pro-Code, CyTOF enables simultaneous detection of dozens of proteins on a single cell, and thus, Pro-Code/CRISPR screens enable high-dimensional phenotyping.

We set out to identify potential regulatory connections between different oncogenes and tumor suppressors with the immunophenotype of CRC cells. To carry out this investigation at a greater scale, we employed the Pro-Code technology. Using a Pro-Code/CRISPR library targeting 34 different commonly mutated genes in CRC, we identified P53 as a positive regulator of CD80. Trp53 deletion led to CD80 loss, causing decreased T cell activation and escape from cancer antigen-specific T cell killing. We found that TP53 binds to the CD80 promoter and directly regulates its transcription. Furthermore, we found that CD80 regulation by TP53 is a conserved program that exists in various cancer cells, normal epithelial cells, as well as macrophages.

## RESULTS

### Establishment of Pro-Code/CRISPR screen in colorectal cancer cells

We sought to understand how different genes associated with cancer cell biology might influence some key immunoregulatory genes in CRC cells. As a model of CRC, we used the mouse CRC cell line CT26 since it is well characterized, extensively used in the literature, and models microsatellite stable (MSS)/non-hypermutated grade IV carcinoma. CT26 cells harbor homozygous Kras-G12D mutation and homozygous deletion of Cdkn2a while having wildtype Trp53^11,12^. We selected the most commonly mutated 34 genes in CRC based on The Cancer Genome Atlas (TCGA) database^13^ and verified that each gene was expressed in CT26 cells by RNA-seq (**Supplementary Table 1**).

To knockout each of the CRC-associated genes, we generated a Pro-Code/CRISPR library encoding CRISPR gRNAs targeting all 34 genes, with 3 gRNAs/gene (102 vectors + 2 control gRNAs). To improve cell coverage in each analysis, we divided the library into two 52 Pro-Code/CRISPR libraries which also allowed for the use of the same unique Pro-Codes for 2 different gene targets. We first wanted to determine whether the Pro-Codes could be detected in CT26 and the distribution of each Pro-Code/CRISPR vector in the library. To avoid the alterations in cell fitness that may be caused by disrupting some of the target genes, we initially used CT26 cells without Cas9 protein, and thus in which the gRNAs would not mediate gene knockout. We transduced CT26 cells at a low multiplicity of infection (MOI) with the 2 libraries of 52 Pro-Code/CRISPR vectors (104 Pro-Code/CRISPR vectors total). After 8 days, cells were stained with metal conjugated antibodies specific for each of the 8 Pro-Code epitopes plus NGFR, which labels all cells with Pro-Codes, and analyzed using CyTOF. We were able to detect all 8 Pro-Code epitopes with a high signal to background separation (**Fig. S1a**). We used a debarcoder software to determine the frequency of each Pro-Code population. Unbiased clustering by viSNE resolved all 104 Pro-Code-expressing cell populations, with each population corresponding to a unique Pro-Code (i.e. exclusively expressing 3 and only 3 epitopes) (**Fig. S1b and S1c**).

Having established we could resolve all 104 Pro-Code/CRISPR CT26 populations by CyTOF, we sought to determine how knockout of the selected genes would impact CT26 fitness. We engineered CT26 to stably express Cas9 and transduced the cells with our Pro-Code/CRISPR library at low MOI. In parallel, we transduced the parent CT26 cells, which do not express Cas9, with the same Pro-Code/CRISPR library, to serve as a control population. We passaged the cells for 14 days to allow ample time for cell population competition and then stained them for the Pro-Code epitopes and analyzed them by CyTOF to determine Pro-Code/CRISPR frequency. While the frequency of most Pro-Code/CRISPR populations did not change more than 2-fold in the CT26-Cas9 compared to the CT26 cells, cells encoding each of the Pro-Code/CRISPR targeting the oncogenes Myc and Kras decreased on average by 4- and 13-fold, respectively (**Fig. S2**). CT26 has Kras-G12D oncogenic mutation and high expression of the Myc oncogene^11^. The loss of cells carrying Pro-Code/CRISPR targeting Kras and Myc fits with the known role of these genes in driving cell proliferation and indicates a non-redundant role in CT26. We also observed enrichment of cells carrying the Pro-Code/CRISPRs targeting Trp53, which encodes the tumor suppressor P53. These findings functionally validated that the oncogenes KRAS and MYC and the tumor suppressor TP53 significantly impact the fitness of CT26 cells.

### Pro-Code/CRISPR screen identifies Kras, Myc, and Trp53 as regulators of CRC immunophenotype

We selected 8 proteins for our immunophenotypic analysis based on their relevance to immune modulation. In addition to MHC-I and PD-L1, our analysis included CD106 (VCAM-1), which is aberrantly expressed in some cancers and can recruit tumor-associated myeloid cells while decreasing CD8 T cell infiltration^14^; CD44, which is a marker for cancer stem cells that evade immune surveillance^15^, CD47 which is a molecule recognized by macrophages as a phagocytosis inhibitor^2^; CD63, which is a marker for exosomes that can regulate tumor immunity^16,17^; CD120b (TNFR2), which forms a heterodimer with CD120a to recognize tumor necrosis factor alpha (TNFα)^18^; and CD80, which is a co-stimulatory molecule that binds to CD28 or CTLA-4 on T-cells to activate or inhibit T-cells, respectively^19^. Examination of the CT26 transcriptome found that each of these genes is expressed at varying levels in steady-state (**Supplementary Table 2**).

CT26 and CT26-Cas9 cells were transduced with the 104 Pro-Code/CRISPR vectors (as two 52 library pools) targeting the 34 prioritized genes. Because some phenotypic markers, such as PD-L1, are upregulated by IFN-γ signaling, we included an arm of cells treated with murine IFN-γ (100 ng/ml) for 16-24 hours before sample analysis. We collected the cells 8 days after library transduction and stained them for the Pro-Code epitopes, MHC-I, CD106, CD80, CD47, PD-L1, CD44, CD63, and CD120b, and analyzed by CyTOF.

We observed that the library transduced CT26 cells with no Cas9 had all 8 immunological effector proteins on the cell surface and that the CT26 upregulated MHC-I and PD-L1 in response to IFN-γ stimulation (**Fig. S3**). Although the expression of most molecules was similar in the majority of Pro-Code/CRISPR populations in the Cas9 expressing cells, there were notable exceptions (**Fig. 1b and 1c; and Fig. S3)**. For example, cells carrying Pro-Code/CRISPR targeting Myc had a marginal increase in MHC-I and an almost 2-fold increase of PD-L1 on the cell surface. MYC has previously been identified as a positive regulator of PD-L1^8^, but our results suggest that in CT26 it may be a negative regulator. Cells encoding Pro-Code/CRISPR targeting Kras had a decrease in the levels of CD106 and this decrease was more prominent in the presence of IFN-γ stimulation. Most strikingly, we observed that cells encoding all three Pro-Code/CRISPR targeting Trp53 (and thus likely p53 KO) had a substantial, sometimes near-complete, decrease in CD80 expression. This decrease was consistent in the presence or absence of IFN-γ and was downregulated to similar levels in all 3 gRNA cell populations. These findings suggest that p53 has a role in controlling the expression of CD80.

**Figure 1.**
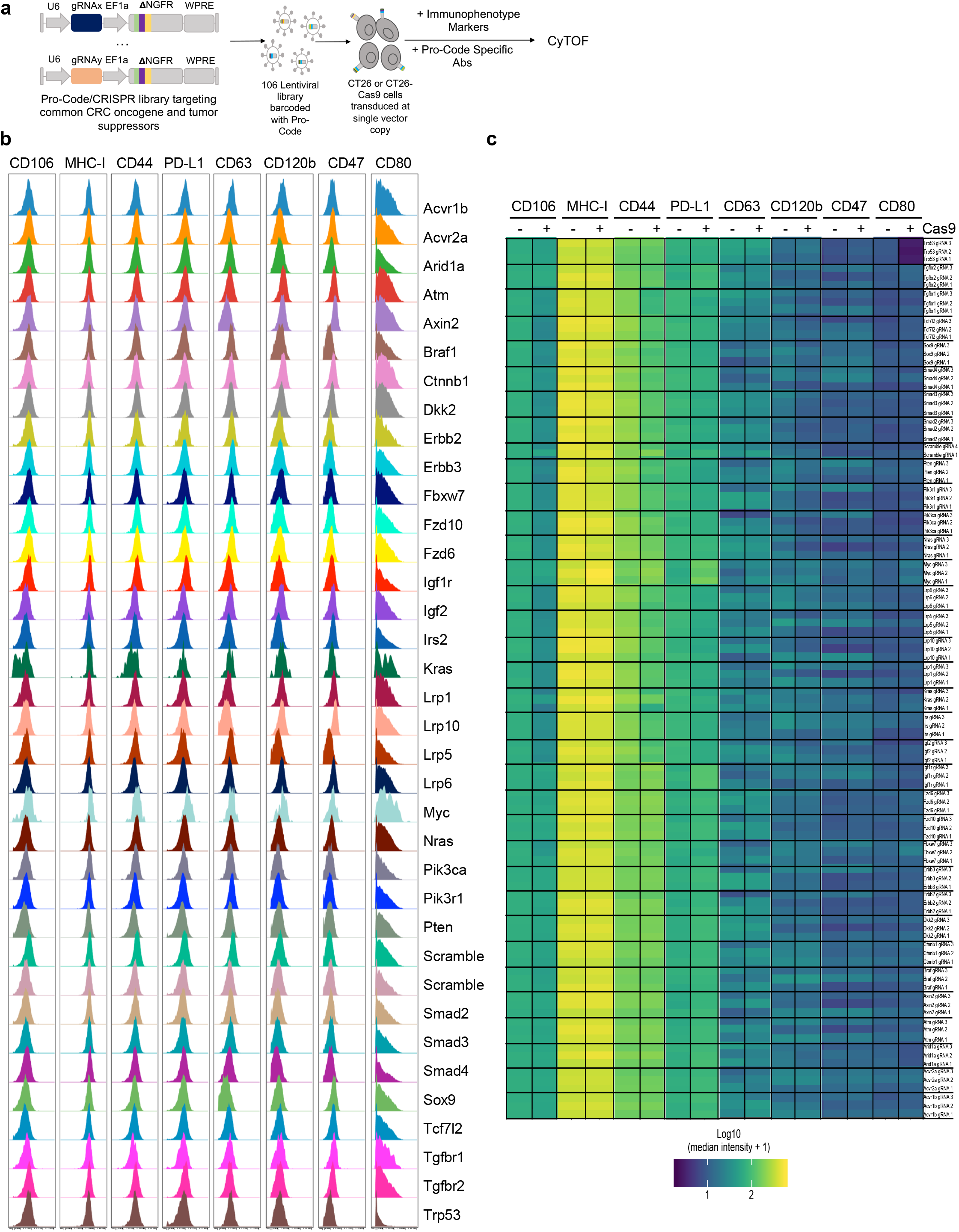
Pro-Code/CRISPR screen of commonly mutated genes in colorectal cancer cells identifies multiple regulators of immunophenotype. a) Schematic of the Pro-Code/CRISPR library, transduction, and phenotypic analysis. CT26 and CT26-Cas9 cells were transduced with 104 Pro-Code/CRISPR vectors, stained, and analyzed by CyTOF mass cytometry. b) Expression of the indicated proteins on each Pro-Code/CRISPR cell population after IFNγ treatment. Shown are representative histograms for each Pro-Code population. The y axis represents cell count normalized by the protein detection channel. c) Heatmap representation of the relative expression of molecules MHC-I, CD106, CD80, CD47, PD-L1, CD44, CD63, and CD120b across all Pro-Code/CRISPR populations after IFNγ treatment. All data is representative of 3 independent experiments.

### P53 is a positive regulator of CD80

To confirm our findings implicating p53 as a regulator of CD80, we transduced CT26 and CT26-Cas9 cells with all three CRISPR gRNAs targeting p53. We stained the cells for CD80 and performed flow cytometry to measure CD80 levels. Impressively, there was a 5-fold decrease in CD80 protein levels in the CT26-Cas9 cells compared to CT26 cells (**Fig. 2a and S4b**).

**Figure 2.**
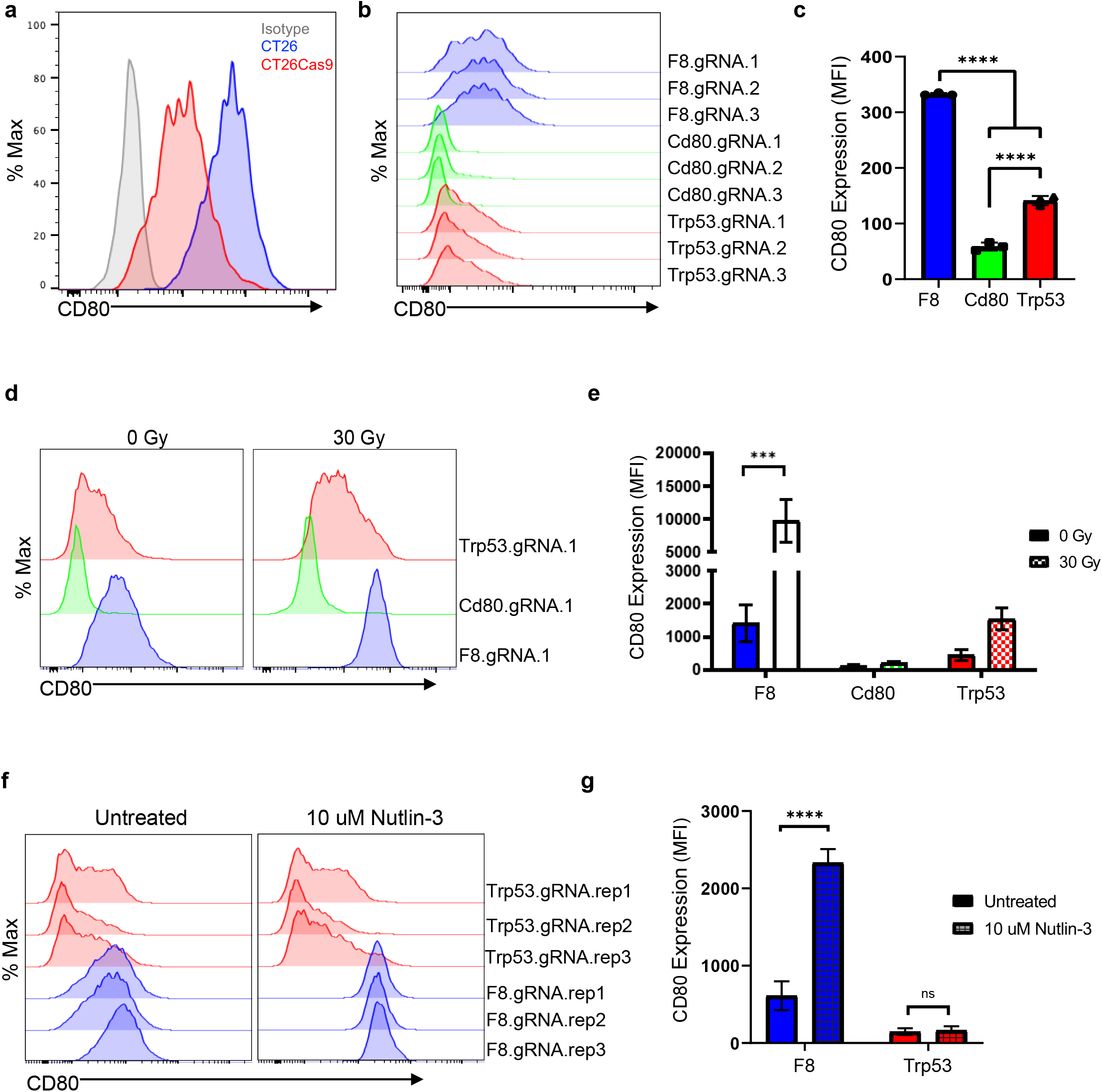
CRISPR/Cas9 knockout of Trp53 reduces the expression of CD80. a) Cell surface CD80 protein levels in CT26 and CT26-Cas9 cells transduced with a pooled library of 3 unique Trp53 Pro-Code/CRISPR vectors; representative histogram is shown. The y-axis represents relative cell count. b) Cell surface CD80 protein levels in CT26-Cas9 cells transduced individually with 3 unique Trp53 gRNA, CD80 gRNA, or control (F8) gRNA vectors; representative histogram shown, the y-axis represents relative cell count. c) Bar graph quantification of (a) as the geometric mean of fluorescent intensity for CD80 (n = 3). Error bars indicate mean ± SD with One-way ANOVA with Tukey’s multiple comparisons, ****p<0.0001. d) CT26-Cas9-GFP cells transduced individually with Trp53 gRNA.1, CD80 gRNA.1, or F8 gRNA.1 vector were either untreated (0 Gy) or gamma-irradiated (30 Gy) and analyzed by flow cytometry. Representative histograms are shown. e) Bar graph quantification of (d) as geometric mean of fluorescent intensity for CD80 (n = 3). Error bars indicate mean ± SEM with Two-way ANOVA with Sidak’s multiple comparisons test, ***p<0.001. f) CT26-Cas9-GFP cells transduced individually with Trp53 gRNA.1, Cd80 gRNA.1, or F8 gRNA.1 vector were either untreated or treated with 10 uM Nutlin-3 and analyzed by flow cytometry. Representative histograms are shown of n=3 biological replicates. g) Bar graph quantification of (f) as the geometric mean of fluorescent intensity for CD80 (n = 3). Error bars indicate mean ± SD with Two-way ANOVA with Tukey’s multiple comparisons test, ****p<0.0001, ns = not significant.

To further validate these findings, we transduced CT26-Cas9 cells with single gRNA targeting Trp53, Cd80, or Factor VIII (F8). F8 is a coagulation co-factor. It is not expressed by CT26 cells and thus serves as a negative control. We carried out a Surveyor Assay of the Trp53 locus which confirmed that the Trp53 gene was efficiently edited by the Trp53 gRNA (**Fig. S4b**). This was supported functionally by the fact that the Trp53 cells had a significant fitness advantage over wildtype CT26 or CT26 that received other gRNA (**Fig. S2a**) which likely leads to the Trp53 knockout cells being highly enriched in the cultures. As expected, CD80 did not change on cells transduced with the F8 gRNAs and was almost completely lost on cells with Cd80 gRNAs. Notably, cells transduced with all three Trp53 gRNAs had a significant decrease in CD80 (**Fig. 2b and 2c**). Thus, these results provide clear validation that P53 is a major regulator of CD80 levels on CT26.

Under homeostatic conditions, P53 can be activated by various insults that cause DNA damage in cells. This typically leads to P53 translocating from the cytoplasm to the nucleus to trigger the expression of necessary genes to allow repair of the damaged genome. One of the many insults that can trigger P53 is ionizing radiation^20^. Owing to this, we tested whether inducing DNA damage in CT26 cells via gamma radiation, a type of ionizing radiation, would alter the expression of CD80 via P53. 30 Gy of radiation altered the morphology of all samples causing the cells to become larger and more granular compared to the non-irradiated samples without causing an increase in dead cells (**Fig. S4c**). We observed that irradiated control F8 knockout cells had on average a 7-fold upregulation of CD80 cell surface levels compared to the unirradiated F8 knockout cells, whereas there was no appreciable difference in CD80 expression between the irradiated and unirradiated Cd80 knockout cells (**Fig. 2d and 2e**). Importantly, CD80 upregulation upon irradiation was significantly diminished in Trp53 knockout cells (**Fig. 2d and 2e**).

To further validate our findings, we employed chemical activation of P53 as an alternative to irradiation. We chose Nutlin-3, which is a specific inhibitor of MDM2, a negative regulator of P53 that functions by targeting TP53 for protein degradation^21^. Similar to irradiation, Nutlin-3 treatment significantly increased CD80 levels in F8 knockout cells and failed to do so in Trp53 knockout cells (**Fig. 2f and 2g**). Thus, whereas loss of Trp53 results in the reduction of CD80, activation of P53 promotes CD80 upregulation. Together these findings indicate that P53 is a positive regulator of CD80.

### TP53 directly regulates CD80 at the transcriptional level

Next, we sought to understand whether the regulation of cell surface CD80 protein levels by P53 was through regulation of CD80 transcription or another mechanism, such as promoting protein stability or translocation. To do this, we examined the expression of Cd80 mRNA in the different cell and treatment groups. We first measured mRNA levels of the canonical transcriptional target of TP53, Cdkn1a (P21)^22^. As expected, irradiation consistently upregulated Cdkn1a mRNA levels in both F8 and Cd80 knockout cells whereas there was no change in Cdkn1a mRNA levels in Trp53 knockout cells (**Fig. 3a**). These observations further validated the functional loss of TP53 in our Trp53 knockout cells. Cd80 expression levels were significantly upregulated upon irradiation of F8 knockout cells while this upregulation was reduced in both Cd80 and Trp53 knockout cells (**Fig. 3a**). These findings suggest that CD80 regulation by P53 occurs at the RNA level.

**Figure 3.**
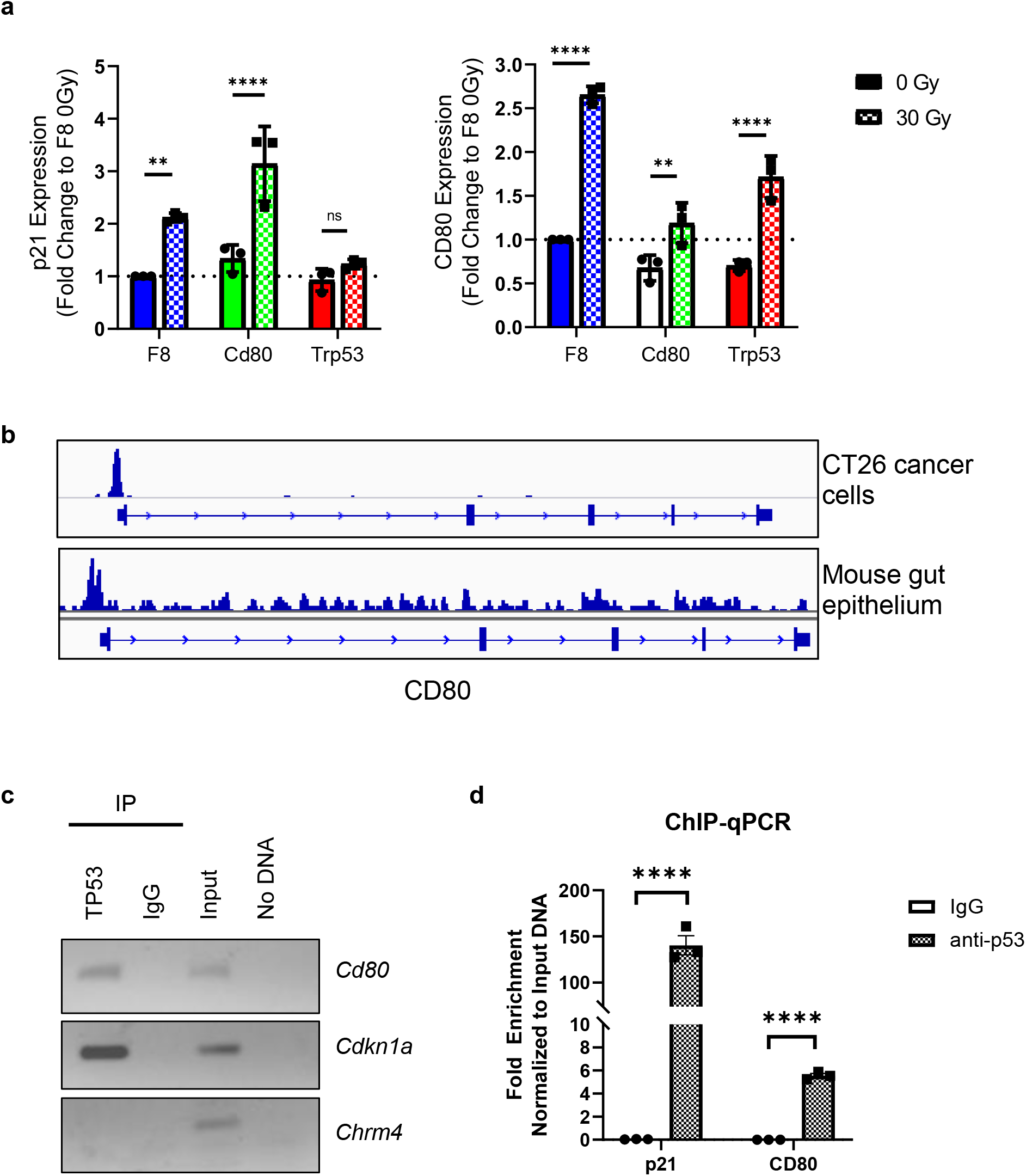
P53 directly regulates Cd80 expression. a) RT-qPCR analysis of Cdkn1a and Cd80 gene expression levels in CT26 cells with CRISPR knockout of either Trp53, or Cd80, or control (F8) with or without gamma irradiation. The graph presents the mean ± SD of the fold change (n = 3). Beta-actin was used to calibrate, and untreated F8 was used to normalize. b) IGV browser visualization of P53 chromatin immunoprecipitation in CT26 colorectal cancer cell line and normal mouse gut epithelium. c) ChIP-qPCR analysis of P53 chromatin immunoprecipitation in CT26 cells for Cdkn1a (positive control), Cd80, and Chrm4 (negative control). ChIP with anti-IgG antibody serves as a negative control. d) Quantification of (c). The graph presents the mean ± SD of the fold change (n = 3). Input DNA was used to normalize. Multiple t-tests, ****p<0.0001.

Since P53 is a transcription factor, it may be a direct regulator of Cd80, though it may also regulate Cd80 indirectly through the regulation of another transcription factor. To address this, we performed a chromatin immunoprecipitation sequencing (ChIP-Seq) experiment with CT26 cells. We identified a TP53 binding site (**Fig. 3b**) with a conserved TP53 binding motif (**Fig. S5a**) at the promoter of CD80. We asked whether CD80 transcriptional regulation by TP53 is an aberration in cancer cells or the same mechanism may also be functional in non-cancerous cells. We repeated TP53 ChIP-Seq with normal mouse gut epithelium and found that indeed TP53 binds to the Cd80 promoter in normal cells (**Fig. 3b**). We further validated these findings by ChIP-PCR and ChIP-qPCR with CT26 cells and we observed a significant enrichment of the Cd80 promoter region in TP53-bound DNA segments (**Fig. 3c and 3d**).

Also, we analyzed publicly available ChIP-Seq data for TP53 and found that TP53 binding to CD80 promoter is conserved across various cell types, including SJSA-1, a human osteosarcoma cell line^23^, and human blood lymphocytes from healthy donors^24^ (**Fig. S5b**). Together these data indicate that TP53 directly binds to the Cd80 promoter and positively regulates its expression.

### TP53 loss enables cancer cells to escape from antigen-specific T cell killing via CD80 downregulation

In homeostatic conditions, CD80 is predominantly expressed by antigen-presenting cells (APC) and its role in T cell activation is well established^25^. CD80 expression is found in some cancer cells, including CRC lines, and overexpression of CD80 in cancer cells stimulates an antitumor immune response^26–28^. To test whether P53 control of CD80 influences T cell priming by CT26 cells, we employed a co-culture model with GFP-specific Jedi T cells^29^. We engineered the F8, Cd80, and Trp53 knockout CT26 cells to stably express GFP as a model antigen. The cells were then co-cultured with naïve Jedi T cells loaded with CellTrace Violet dye. After 5 days of co-culture, the amount of dye dilution in the Jedi T cells was analyzed using flow cytometry (**Fig. 4a**). Whereas control CT26-GFP cells (F8 gRNA) were able to prime and promote the proliferation of Jedi T cells, there was a significant reduction in priming mediated by the Cd80 and Trp53 knockouts cells, as indicated by the number of Jedi T cells that were not induced to divide (**Fig. 4b and 4c**). This indicates that P53-mediated control of CD80 levels is functionally consequent to enable CT26 priming of naïve antigen-specific T-cells.

**Figure 4.**
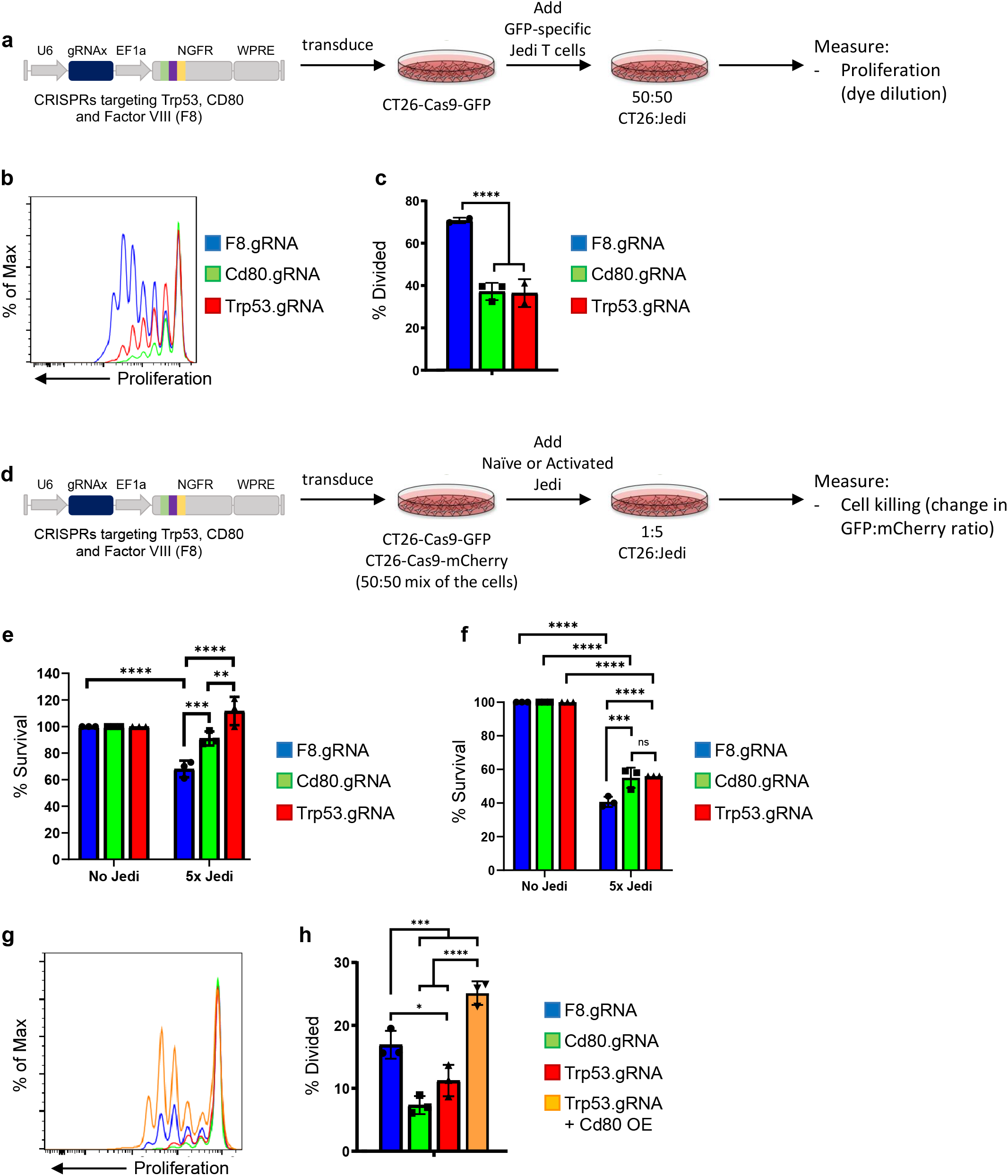
Reduced CD80 upon loss of P53 in cancer cells decreases activation of naïve tumor antigen-specific T cells. a) Schematic of the T cell co-culture activation experiment. b) The proliferation of GFP-specific CD8+ T-cells (Jedi T-cells). Naïve Jedi T-cells were stained with CellTrace violet dye then co-cultured with CT26-Cas9-GFP cells for 5 days in the presence IFNγ and analyzed via flow cytometry. Representative histograms are shown. c) Quantification of proliferating naïve Jedi T-cells. Bar plots represent mean ± SD of the percentage of dividing cells in naïve Jedi cells co-cultured with Trp53 gRNA, CD80 gRNA, or F8 gRNA transduced CT26-Cas9-GFP cells (n = 3). Two-way ANOVA with Sidak’s multiple comparisons test ****p < 0.0001, ns = not significant d) Schematic of co-culture killing experiment. e) The frequency of GFP+ CT26-Cas9 cells (transduced with p53, CD80, or F8 gRNAs) after co-culture with naïve Jedi T cells (1:5 ratio) (as described in D). Bar graphs present the mean ± SEM (n = 3). Two-way ANOVA with Sidak’s multiple comparisons, ****p<0.0001, ***p<0.001, **p<0.01, *p<0.05, ns = not significant. f) The frequency of GFP+ CT26-Cas9 cells (transduced with p53, CD80, or F8 gRNAs) after co-culture with activated Jedi T cells (activated with anti-CD28, anti-CD3 and IL-2). Bar graphs present the mean ± SEM (n = 3). Two-way ANOVA with Sidak’s multiple comparisons, ****p<0.0001, ***p<0.001, **p<0.01, *p<0.05, ns = not significant. g) The proliferation of GFP-specific CD8+ T cells (Jedi T cells). Naïve Jedi T cells were stained with CellTrace violet dye then co-cultured with Trp53 gRNA, CD80 gRNA, or F8 gRNA with CD80 overexpressed transduced CT26-Cas9-GFP cells for 5 days with IFNγ and analyzed via flow cytometry. Representative histograms are shown. h) Quantification of proliferating naïve Jedi T-cells. Bar plots represent mean ± SD of the percentage of dividing cells in naïve Jedi cells co-cultured as described in (I) (n = 3). Two-way ANOVA with Sidak’s multiple comparisons test, ****p < 0.0001, ***p<0.001, **p<0.01, *p<0.05.

We next evaluated whether the primed T cells were functional. CT26 cells with individual knockouts of F8, Cd80, or Trp53 were transduced to stably overexpress either GFP, the antigenic target of Jedi T cells, or an unrelated antigen, mCherry. In each condition, equal numbers of GFP and mCherry expressing cells were mixed and then co-cultured with naïve Jedi T cells for 5 days. Antigen-specific cancer cell killing was measured as a ratio of GFP+/mCherry+ target cells by flow cytometry (**Fig. 4d**). In the control cultures, there was a 40% reduction in the ratio of GFP/mCherry+ cells, indicative of Jedi T cell killing of their antigen-expressing target cells. In contrast, in the cultures in which Cd80 and Trp53 were knocked out, there was little to no reduction in the GFP/RFP+ cell ratio, indicating the GFP+ cells were not being killed by the Jedi T cells (**Fig. 4e**). This is consistent with the lack of priming of the Jedi T cells when Cd80 or Trp53 are knocked out. To see if strong activation of the T cells could surmount this defect, we activated the Jedi with anti-CD3/anti-CD28 beads, and then performed the co-culture. As expected, target cell killing was greater in all cultures (as the activation circumvented the need for priming by the CT26) though there was still a small, yet significant difference in the level of GFP+ cell killing in the control culture compared to the Cd80 and Trp53 knockout cultures (**Fig. 4f**).

To test whether decreased T cell priming and activation upon TP53 loss in cancer cells were due to downregulated CD80, we overexpressed mouse Cd80 in CT26 cells with Trp53 knockout (**Fig. S6**). We then repeated antigen-specific T cell priming experiments as described above with the addition of the CD80 rescue cell line. We found CD80 overexpression in CT26-GFP cells with Trp53 knockout rescued Jedi T cell priming and proliferation (**Fig. 4g and 4h**). Together, these data demonstrate that CD80 expressed on cancer cells promotes both antigen-specific T cell priming and activation and that loss of TP53 abrogates this activation and helps enable escape from killing by naïve T cells.

### P53 regulates CD80 in macrophages

CD80 is most predominantly expressed by professional APCs such as monocytes and macrophages where it plays a key functional role in the APC activity of these cells. To determine if P53 can also regulate CD80 in macrophages, we generated mouse bone marrow-derived macrophages and treated them with Nutlin-3 or radiation to induce P53 signaling. As a positive control, we treated the cells with LPS, and, as expected, this induced high levels of CD80, as well as PD-L1, another LPS-induced gene. Notably, TP53 activation by either irradiation or Nutlin-3 treatment did not upregulate PD-L1 but was sufficient to upregulate CD80 (**Fig. 5a and 5b**; 900 vs 1,450 CD80 MFI in DMSO vs Nutlin). To rule out any potential CD80 induction by cell death upon irradiation or Nutlin-3 treatment, we also co-cultured the macrophages with irradiated monocytes and macrophages, which did not induce CD80 (**Fig. 5a and 5b**). We also analyzed a publicly available gene expression dataset of human monocyte-derived macrophages which were treated with DMSO, LPS, and Nutlin-324. Similar to our mouse macrophage cultures, CD80 gene expression was upregulated upon Nutlin-3 treatment (**Fig. S7a**).

**Figure 5.**
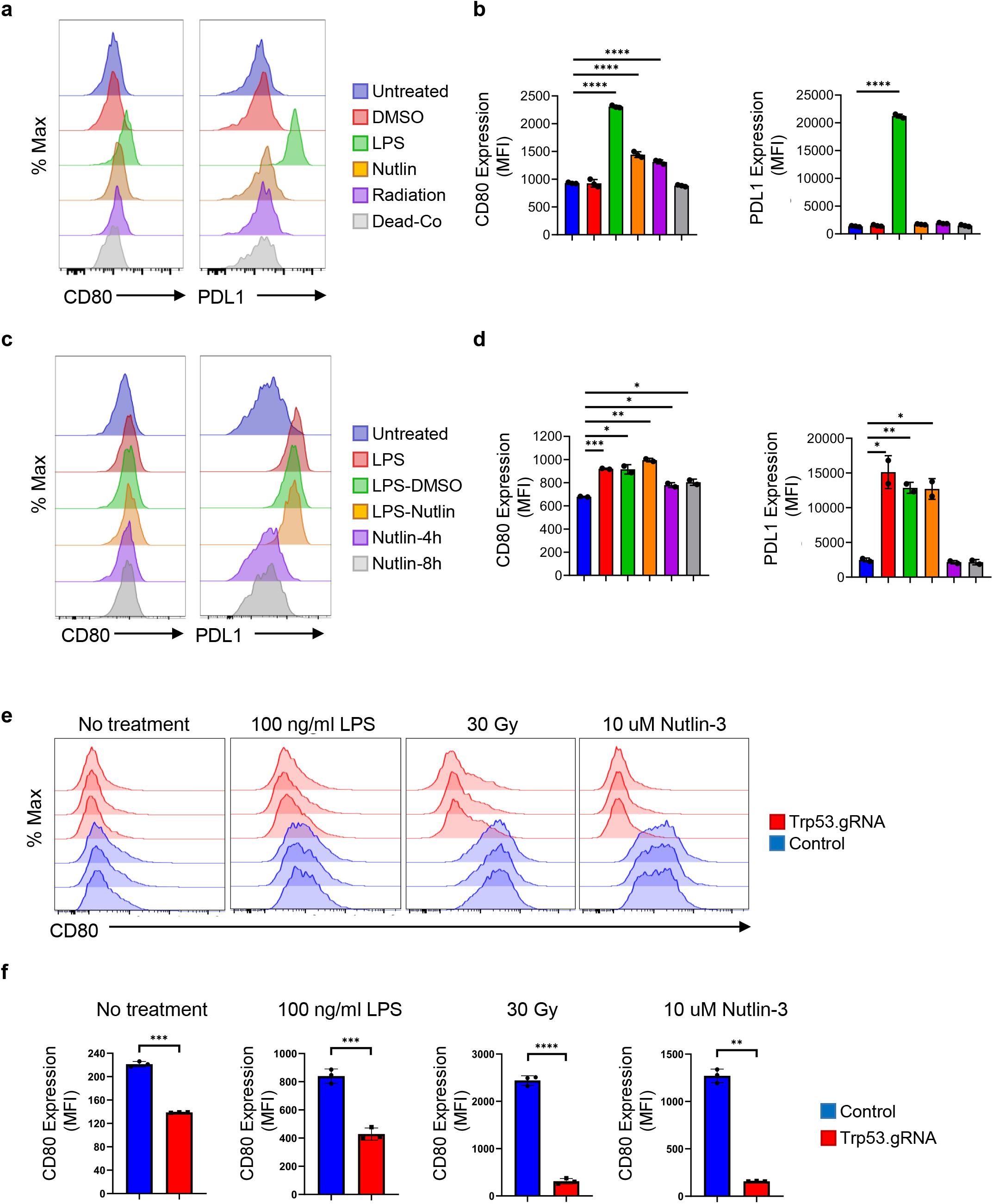
P53 regulates the co-stimulatory molecule CD80 in macrophages. a) Flow cytometry histograms comparing cell surface CD80 and PD-L1 levels in mouse bone marrow-derived macrophages untreated or treated with either DMSO, LPS (100 ng/ml), Nutlin-3 (10 µM), or radiation (10 Gy), or co-cultured with irradiated apoptotic cells (dead-co) for 24 hours. b) Bar graph quantification of (a) as geometric mean of fluorescent intensity for CD80 or PD-L1 (n = 3). Error bars indicate mean ± SD with unpaired t tests, ****p < 0.0001, ***p < 0.001. c) Flow cytometry histograms comparing cell surface CD80 and PD-L1 levels in mouse tumor-associated macrophages isolated from ascites of ID8 ovarian tumor-bearing mice and treated as indicated ex vivo. LPS alone is for 8 hours while LPS-DMSO and LPS-Nutlin treatments are for 4 hours for each consecutive treatment. The drug concentrations are the same as in (a). d) Bar graph quantification of (c) as geometric mean of fluorescent intensity for CD80 or PD-L1 (n = 2 mice). Error bars indicate mean ± SD with unpaired t tests, ****p < 0.0001, ***p < 0.001. e) Raw264.7 macrophage cells engineered to stably express Cas9 were transduced individually with 3 unique Trp53 gRNA vectors. Cell surface CD80 levels were analyzed by flow cytometry at baseline and after treatment with either LPS (100 ng/ml), Nutlin-3 (10 µM), or irradiation (30 Gy). Representative histograms are shown. f) Bar graph quantification of (e) as geometric mean of fluorescent intensity for CD80 (n = 3). Error bars indicate mean ± SD with unpaired t tests, ****p < 0.0001, ***p < 0.001, **p<0.01.

Macrophages constitute a considerable fraction of the tumor microenvironment in many solid tumors and are hijacked by cancer cells to support tumor growth^30^. To test whether P53 regulation of CD80 is conserved in tumor-associated macrophages (TAMs), we isolated ascites from an orthotopic mouse model of ovarian cancer (ID8) and enriched TAMs ex vivo. Similar to bone marrow-derived macrophages, when we treated TAMs with Nutlin-3 to stabilize P53, there was an upregulation of CD80, but not PD-L1 (**Fig. 5c and 5d**).

For genetic perturbation studies in macrophages, we generated F8, Cd80, or Trp53 knockouts using Raw264.7 mouse macrophage cells by CRISPR/Cas9. We then treated the cells with LPS, Nutlin-3, or radiation. Already in steady-state TP53 loss significantly decreased baseline CD80 levels in the macrophages. Similar to our findings in cancer cells, bone marrow-derived macrophages, and TAMs, TP53 signaling, induced by either Nutlin-3 treatment or irradiation, significantly upregulated CD80 protein levels, which was abrogated in Trp53 deleted cells (**Fig. 5e and 5f**).

Hepatic stellate cells are liver-resident cells that help form the stroma of the tissue^31^. Stellate cells do not express Cd80 in steady-state (**Fig. S7b**) but we wondered if P53 activation might induce its upregulation in these cells. To test this, we used JS1 cells, a mouse hepatic stellate cell line. We generated Cd80 or Trp53 knockouts using CRISPR/Cas9 or introduced the control F8 gRNA. The cells were either untreated (0 Gy) or irradiated (30 Gy) and 16-24 hours later CD80 expression was analyzed by flow cytometry. JS1 cells did not have any detectable cell surface CD80 in the baseline but irradiation induced CD80 expression on a subset of the control cells. In contrast, irradiation did not induce CD80 upregulation in stellate cells in which Trp53 or Cd80 were knocked out (**Fig. S7c**). These data extend the finding that TP53 is a positive regulator of CD80 to non-malignant cells, including hepatic stellate cells and macrophages.

## DISCUSSION

Using Pro-Code/CRISPR genomics, we screened common oncogenes and tumor suppressors to determine if they have a role in regulating the immunophenotype of CRC cells. Analysis of eight cell surface proteins in CT26 CRC cells with 104 different CRISPRs targeting 34 genes found that VCAM-1, PD-L1, and CD80 are regulated by KRAS, MYC, and P53, respectively. Genetic, pharmaceutical, and physical perturbations that either inhibited or activated P53 confirmed that P53 positively regulates CD80. Gene expression and chromatin immunoprecipitation experiments showed that this regulation is direct and at the transcriptional level. Functionally, modulation of CD80 by P53 in cancer cells altered antigen-specific cytotoxic T cell response against the cancer cells. We found that the P53 – CD80 signaling axis is preserved in various cell types, including macrophages.

In response to various cellular stress, most notably DNA damage, stabilized P53 regulates transcription of hundreds of genes to orchestrate a cellular response that may include cell cycle arrest, DNA repair, senescence, or apoptosis^32^. As the most frequently mutated gene in cancer, TP53 has been extensively studied in various contexts. However, its role in controlling the immunophenotype of cells is less well characterized. Muñoz-Fontela and colleagues first showed that TP53 functions in a positive feedback loop in the type I interferon response^33^ and other studies have reported P53 target genes as having roles within different innate immune pathways^34^. TP53 directly regulates the transcription of FOXP3 thereby induces the regulatory T cell program in CD4 T cells ^35^. TP53 activation also shifts macrophage functional state from immunosuppressive M2 towards pro-inflammatory M1 state^36^. In addition, P53 has been reported to positively regulates MHC-I antigen presentation by transcriptional modulation of TAP1 and ERAP1^37,38^. P53 promotes transcription of miR-34a, which suppresses PDL1^39^. Another cancer cell-intrinsic immunomodulatory role of P53 is the regulation of various Toll-like receptors, such as TLR3 and TLR5^40^.

In line with these mechanisms where wild-type P53 activity increases immune recognition of cells, our findings indicate that P53 also upregulates CD80, a key co-stimulatory molecule. The activation of a naïve antigen-specific T cell requires two signals, signal 1 is provided by peptide/MHC, and signal 2 is provided by CD80 or CD86, which bind to CD28 on T cells to initiate a signaling cascade that enables a naïve T cell to be activated. T cells also express on their surface CTLA4, a negative regulator of T cell activation which competes with CD28 for CD80/86 binding^19^. Low levels of cell surface CD80 can promote immune evasion by increasing CTLA-4 mediated inhibitory signaling in T cells due to the higher binding affinity of CD80 to CTLA-4 compared to its affinity for CD28^28^.

CD80 is upregulated in human preneoplastic colon epithelial cells, and blockage or loss of CD80 enhances dysplasia development in mouse colon^41^. The Melero lab previously reported that CD80 is expressed at relatively low levels by CT26 CRC cells and that it serves as an immune evasive purpose, as knockdown of CD80 by RNAi led to tumor clearance in immunocompetent mice but not in immune-deficient mice^28^. We did not carry out in vivo studies since the role of CD80 in CT26 tumor engraftment was demonstrated in this previous work. Instead, our focus here was on establishing one of the regulators of CD80 in these cells, which appears to be P53. Future studies will be helpful to better assess the in vivo relevance of the P53/CD80 axis, though the pleiotropic functions of P53 make it more challenging to establish discrete functions related to a single molecule in vivo.

CD80 is most abundantly expressed by myeloid cells, predominately APCs, and the canonical role of CD80 is in providing a crucial signal to T cells in APC – T cell synapses^25^. While the role of CD80 in APCs is well established, the immunomodulatory role of P53 in these cells is less known. Previous studies found that P53 upregulates many proinflammatory cytokines in human macrophages and that P53 activity can limit M2 polarization of macrophages^24,36^. Our findings indicate that P53 can also positively regulate CD80 expression on macrophages. It may be that CD80 upregulation by P53 helps to enable APCs to activate a T cell response as a mechanism of promoting immune editing of cells with DNA damage. This will be something to further investigate, particularly in the context of APCs that are exposed to genotoxic stresses, such as Langerhans’s cells in the skin or tumors undergoing radiotherapy.

## ACKNOWLEDGMENTS

We thank A. Rodriguez for helpful discussions. We also thank the Human Immune Monitoring Center and the Flow Cytometry Core for technical assistance. B.D.B. was supported by the National Institutes of Health (NIH) R33CA223947 and R01AT011326 and an award from the Cancer Research Institute (CRI). E.Z. was supported by R01DK106593. G.M. was supported by the NIH K00CA223015 from the National Cancer Institute.

## AUTHOR CONTRIBUTIONS

E.A. and G.M. designed and performed experiments, analyzed data, and wrote the manuscript. M.D. designed experiments and analyzed data. R.Z. performed experiments and analyzed data. S.R. analyzed data. A.B. performed experiments, analyzed data, and proofed the manuscript. R.P. analyzed data. B.D.B. designed and supervised the research, analyzed data, and wrote the manuscript.

## MATERIALS AND METHODS

### Mice

C57BL/6 mice were purchased from Jackson Laboratory. Jedi mice^29^ were from our own established colonies and are also available from Jackson Laboratory. All mice were housed in an environmentally controlled, pathogen-free facility. At the time of experimentation, mice were 6-8 weeks of age. All mouse experiments were performed in accordance with the Institutional Animal Care and Use Committee of the Icahn School of Medicine at Mount Sinai.

### Cell culture

293T cells were maintained in Isocove’s Modified Dulbecco’s Medium (IMDM; Gibco) supplemented with 100 U penicillin/streptomycin per ml, 2mM L-Glutamine (Gibco), and 10% fetal bovine serum (FBS; Gibco). Cells were passaged up to 20 times (washed with PBS, detached from the plate with 0.05% Trypsin-EDTA (Gibco), and replated). CT26 cells are a BALB/c cell line of colorectal carcinoma. CT26 cells were cultured in RPMI1640 (Gibco) supplemented with 10% heat-inactivated FBS, 100 U/ml penicillin/streptomycin, and 2mM L-Glutamine. Raw264.7 cells, a BALB/c cell line of macrophages, were cultured in DMEM with 10% heat-inactivated FBS (GIBCO), 100 U/ml penicillin/streptomycin (GIBCO), and 2 mM L-Glutamine. Cells were kept at maximum confluency of 70% and passaged up to 20 times as described for 293T cells. 293T and CT26 cell lines were purchased from ATCC and were regularly validated to be free of mycoplasma.

### Cell treatments

The cells were seeded 16-24 hours before any treatment. For radiation, the cells were irradiated once at 30 Gy with an X-ray source (RS 2000). Unless otherwise noted, Nutlin-3 (SigmaAldrich) was used at 10 uM final concentration, and LPS (Invitrogen) was used at 100 ng/ul final concentration.

### Harvesting mouse colon epithelium cells

Mice were sacrificed, and the colon was dissected distal to the cecum and proximal to the rectum. The colon was flushed with ice-cold PBS and split open longitudinally. Epithelial cells were collected using a cell scraper to separate them from the underlying mesenchyme.

### T cell activation assay

CD8+ T cells were isolated from spleens of Jedi mice. Splenic cell suspensions were obtained by mechanical disruption and filtering through a 70-mm cell strainer. Red blood cells were lysed using RBC lysis buffer (eBioscience), and CD8+ T cells were negatively selected using EasySep mouse CD8+ T cells isolation kit (StemCell Technologies), following manufacturer’s instructions. CD8+ cells were stained using CellTrace violet proliferation kit per the manufacturer’s protocol. CT26-Cas9-GFP cells were transduced with either Trp53, F8, or CD80 Pro-Code/CRISPR vectors at an MOI of 7 and cell sorted based on NGFR expression. A 50:50 mix of CT26-Cas9-GFP and naïve CellTrace stained T-cell in 24-well plates (8⨯10^4^ cells per well). For indicated samples, co-cultures were treated with 100 ng/ml mouse recombinant IFNg (Peprotech). Activation of T-cells was assessed by flow cytometry on day 5.

### T cell killing assay

Jedi T cells were isolated as described above. For activated T cell killing, Jedi T cells were activated for 3 days with 5 mg/ml plate-bound anti-CD3 mAb (clone 2C11, BioXCell), 1 mg/ml anti-CD28 mAb (clone 37.51, BioXCell) and 20 ng/ml mouse recombinant IL-2 (Peprotech) in RPMI with 10% FBS, 100 U/ml penicillin/streptomycin, 2 mM L-glutamine, 1% non-essential amino acids, 1 mM sodium pyruvate, 55 mM 2-mercaptoethanol, and 20 mM HEPES, otherwise cells were immediately used for co-culture post isolation. CT26-Cas9-GFP cells were transduced with either Trp53, F8, or CD80 Pro-Code/CRISPR vectors at an MOI of 7 and cell sorted based on NGFR expression. A 50:50 mix of GFP+ (target) and mCherry+ (bystander) CT26-Cas9 cells were plated in 24-well plates (4×10^4^ cells per well). Naïve or activated T-cells were added to the wells 6 hours later, at a ratio of 5:1 (E: T), with 100 ng/ml mouse recombinant IFNg (Peprotech) addition. The killing was assessed by flow cytometry on day 3 for activated T-cell co-cultures and 5 for naïve T-cell co-cultures.

### Macrophage generation/isolation

Bone marrow cells were isolated from the femur and tibia of C57BL/6 mice by flushing the bones with PBS. Cells were pipetted several times to make single-cell suspension and passed through a 70 um cell strainer followed by cold PBS wash and centrifugation at 350 g for 5 minutes. Cells were resuspended, counted, and seeded at 1 × 10^6 cells/ml density in media (RPMI with 10% FBS, 100 U/ml penicillin/streptomycin, 2 mM L-glutamine, 1% non-essential amino acids, 1 mM sodium pyruvate, 55 mM 2-mercaptoethanol, and 20 mM HEPES) supplemented with 10 ng/ml murine M-CSF (Peprotech) in 24-well plate. Non-adherent cells were washed away and the cells were supplemented with fresh medium with M-CSF every other day. By day 6, the macrophages were ready for downstream treatments. Tumor-associated macrophages were isolated from ID8 tumor-bearing mice. 5 × 10^6 ID8-VEGF-Defensin-Brca1-/-cells were transplanted into 6-8 weeks old female C57BL/6 mice by intraperitoneal injection. By 3 weeks post-transplantation, ascites were collected from the peritoneum of tumor-bearing mice by aspiration with syringes. The ascites were then centrifuged at 350 g for 5 minutes and red blood cells were lysed with RBC lysis buffer (eBioscience) followed by PBS wash and centrifugation. The resuspended cells were counted and then seeded at 1⨯ 10^6 cells/ml density in the same macrophage media described above supplemented with 10 ng/ml murine M-CSF in a 24-well plate. The next day, non-adherent cells were washed away and the cells were supplemented with fresh medium with M-CSF before the downstream treatments the same day.

### Vector construction

To clone gRNA sequences, Pro-Code vectors were digested with BbsI and ligated with pairs of annealed oligo sequences (forward oligo design: 50 CACCG(N)20; reverse oligo design: 50 AAAC(N)20C, where (N)20 is the sequence of guide RNA or its reverse complement counterpart). sgRNA sequences (**Supplementary Table 3**) were obtained from Brie (mouse) CRISPR libraries^42^. TOP10 competent cells were used for all subsequent plasmids except lentiCRISPR v2 (Addgene plasmid no. 52961)^43^, which was propagated using NEB stable competent cells (New England BioLabs). All plasmids were purified using the ZR Plasmid Miniprep Classic kit (Zymo Research).

### Lentiviral vector production and titration

Lentiviral vectors were produced as previously described in detail^44^. Briefly, 293T cells were seeded 24 hours before calcium phosphate transfection with third-generation VSV-pseudotyped packaging plasmids and the transfer plasmids. Supernatants were then collected 30 hours after transfection, passed through a 0.22-mm filter, purified by ultracentrifugation, aliquoted, and stored at -80 C. Viral titer was estimated as TU/ml by flow cytometry analysis of NGFR+ cells on 293T cells by limiting dilution. LentiCRISPR v2 transfer plasmid encoding Cas9 transgene and a puromycin resistant cassette was used to generate Cas9 lentivirus. To produce LV Pro-Code libraries, equimolar amounts of single plasmids were pooled and subsequently used for vector production.

### Lentiviral transduction

3⨯10^5^ CT26 cells were seeded in a 6-well plate (5⨯10^4^ cells/well). 24 hours later, cells were transduced with Cas9 lentivirus in the presence of 5 mg/ml polybrene (Millipore). 16 hours after transduction, media was changed to remove the virus. 48 hours after transduction, cells were treated overnight with 10 mg/ml puromycin (ThermoFisher) to remove all non-transduced cells. Puromycin treatment was repeated two additional times to ensure cell purity. For library generation, CT26 cells (+/-Cas9) were seeded 24 hours before transduction at 3⨯10^5^ in a 6-well plate, then transduced at an MOI of 1 (fewer than 10% of cells transduced). 16 hours after transduction, media was changed to remove the virus. 96 hours after transduction, NGFR+ cells were sorted using flow cytometry.

### Flow cytometry

Before FACS analysis, adherent cells were detached with 0.05% trypsin-EDTA, or PBS with 1 mM EDTA for macrophages, washed and resuspended in sterile PBS. For analysis of GFP expression, cells were washed and resuspended in flow buffer (PBS, 2 mM EDTA, 0.5% BSA). For immune staining, flow buffer was supplemented either with anti-mouse CD16/CD32 antibody (eBioscience) or Human TruStain FcX Fc Receptor Blocking Solution (BioLegend). The following antibodies were used for flow analysis: CD271-Alexa Fluor 647 (BD Biosciences 560326), CD271-PE (BD Biosciences 560326), CD11b-PE (eBioscience 12-0112-82), F4/80-Alexa Fluor 488 (Biolegend 123120), F4/80-PerCP-Cyanine5.5 (eBioscience 45-4801-82), CD80-APC (eBioscience 17-0801-82), CD80-PE (eBioscience 12-0801-82), PDL1-PE-Cy7 (eBioscience 25-5982-82). Data were acquired using BD Fortessa (BD) and analysis was performed using FlowJo Software (FlowJo, LLC).

### Mass cytometry

Antibodies were either purchased pre-conjugated from Fluidigm or purchased purified and conjugated in-house using MaxPar X8 Polymer Kits (Fluidigm) according to the manufacturer’s instructions. Following antibodies were used for CyTOF staining: HA tag-147Sm (clone 6E2, Cell Signaling), V5 tag-152Sm (Thermo Fisher Scientific), anti-DYKDDDDK (FLAG) tag-175Lu (clone 5A8E5, GenScript), VSVg tag-158Gd (rabbit pAb, Thermo Fisher Scientific), E tag-154Sm (clone 10B11, Abcam), NWSHPQFEK (NWS) tag-159Tb (clone 5A9F9, GenScript), AU1-162Dy (clone AU1, BioLegend), AU5-169Tm (clone AU5, BioLegend), H2Kd-149Sm (clone SF1-1.1.1, eBioscience), anti-mouse CD274-151Sm (MIH5, eBioscience), anti-mouse CD80-173 (clone 16-10A1, eBioscience), anti-mouse CD47-168Er (clone miap301, BioLegend), anti-mouse CD63-160Gd (clone NVG-2, BioLegend), anti-mouse CD120b-166Er (clone TR75-32.4, BioLegend), anti-mouse CD106-146Nd (clone MVCAM.A, BioLegend), and anti-mouse CD44-150Nd (clone IM7, Fluidigm). Before CyTOF analysis, cells were collected, washed, resuspended in media, and stained for viability with Cell-ID Intercalator-103Rh for 15 min at 37°C. To avoid non-specific staining, cells were subsequently blocked in flow buffer supplemented with either anti-mouse CD16/CD32 antibody (eBioscience) or Human TruStain FcX Fc Receptor Blocking Solution (BioLegend) for 30 min on ice. Next, cells were stained for cell surface antigens, fixed, and permeabilized using BD Cytofix/Cytoperm solution (BD Biosciences) and stained with the tag antibodies for 30 min on ice. After intracellular/tag staining, cells were washed and incubated in 125 nM Ir intercalator (Fluidigm) diluted in PBS containing 2.4% formaldehyde for 30 min at RT, washed and stored in PBS at 4°C. Immediately before the acquisition, samples were washed once with PBS, once with de-ionized water, and then resuspended at a concentration of 10^6^ cells/ml in deionized water containing a 1:20 dilution of Equation 4 Element Beads (Fluidigm). The samples were acquired on either a CyTOF2 or Helios (both Fluidigm) equipped with a SuperSampler fluidics system (Victorian Airships) at an event rate of < 500 events/second at the Human Immune Monitoring Center (HIMC) at Mount Sinai. After the acquisition, the data were normalized using bead-based normalization using the CyTOF software. The data were gated to exclude residual normalization beads, debris, dead cells, and doublets, leaving NGFR+ events for clustering and high dimensional analyses.

### Western blot

For each sample, 10^6^ cells were lysed in RIPA lysis buffer (150 mM sodium chloride, 50 mM Tris, pH 8.0, 1.0% NP-40, 0.5% sodium deoxycholate, 0.1% SDS) supplemented with protease inhibitors (Roche). The amount of protein was determined using QuBit Protein Assay Kit (ThermoFisher) following the manufacturer’s instruction and 80 ug of protein was loaded per well. Western blot was performed according to the supplier’s instructions using rabbit polyclonal anti-p53 antibody (CM5, Leica) and rabbit-monoclonal anti-vinculin antibody (EPR8185, Abcam) for control.

### Surveyor mutation detection assay

Total DNA was extracted from cultured F8.1, Trp53.1, Trp53.2, and Trp53.3 CT26Cas9GFP cells using a DNA extraction kit (Qiagen) per the manufacturer’s direction. The CRISPR targeted region in the p53 locus was PCR amplified with GoTaq Polymerase (ThermoFisher) using the following primers:

Trp53_NHEJ_Forward1: GGAAAGGTCCCAGTCCTCTCTTTGC

Trp53_NHEJ_Reverse1: GGTGAAGTCGCTCCCTACCTCACTA

Trp53_NHEJ_Forward2: GTGAGTGGATCTTTTTGGGGCCCTT

Trp53_NHEJ_Reverse2: AAACTCTGAGGCACAGTCTACAGGC

PCR products were checked via 1% agarose gel electrophoresis for quantity and quality, then hybridized to form heteroduplexes. Surveyor mutation detection (IDT) was performed according to the manufacturer’s protocol. The resulting digested DNA was visualized on 1% agarose gel.

### mRNA sequencing

Total RNA was extracted from cultured CT26 cells using QIAzol lysis reagent (Qiagen) with miRNeasy Mini kit (QIAGEN) following the manufacturer’s instructions. For RNA-seq, RNA integrity and concentration were determined on the Agilent 2100 Bioanalyzer (Agilent; Palo Alto, CA, USA). 20 ng of total RNA was pre-amplified using the Nugen Ovation RNA-seq System 2 (Nugen, San Carlos, CA), and then prepared for sequencing on the Illumina platform using the Tru-seq RNA Library Prep V2 (Illumina, San Diego, CA). The barcoded samples were multiplexed and sequenced on an Illumina NextSeq 550 to a depth of at least 50,000,000 reads per sample. The reads were mapped using RNA Dashboard software (http://katahdin.mssm.edu/html/scripts/resources.pl). Normalized gene expression levels were calculated by RPKM using exon mapping reads.

### mRNA expression analysis

For RT-qPCR, total RNA was extracted from cultured CT26 cells using QIAzol lysis reagent (Qiagen) with miRNeasy Mini kit (Qiagen) following the manufacturer’s instructions. RNA was then reverse-transcribed using an RNA-to-cDNA kit (Applied Biosystems). qPCR was performed using the SYBR green PCR 2x master mix (Thermo Scientific) with the following primers:

Cd80 Forward: AGAAGGAAAGAGGAACGTATGAAG

Cd80 Reverse: TGGGTTTCCAGACTCAGTTATG

Cdkn1a Forward: GTCTTGCACTCTGGTGTCTG

Cdkn1a Reverse: GATAGAAATCTGTCAGGCTGGTC

Actin Forward: CTAAGGCCAACCGTGAAAAG

Actin Reverse: ACCAGAGGCATACAGGGACA

For ChIP-qPCR, qPCR was performed on a sample of ChIP DNA before library prep using SYBR green PCR 2x master mix (Thermo Scientific) with the following primers:

Cd80 Forward: GCCATGAGGGCAACTCAAA

Cd80 Reverse: CCCGCAAGCAGAATCCTTT

Cdkn1a Forward: GCTGGTAGTTGGGTATCATCAG

Cdkn1a Reverse: CTTTCTGGCCTTCAGGAACA

Chrm4 Forward: CCTCCCCCAACTCCACTTTT

Chrm4 Reverse: GGAGGTTGGAGGGAGAAGGA

### Chromatin immunoprecipitation (ChIP)

Double crosslink ChIP was performed as previously described with modifications^45^. Cells were rinsed with PBS and double-crosslinked at room temperature, first using 0.25 M disuccinimidyl glutarate (Thermo Fisher) for 45 minutes followed by 1% formaldehyde for 10 minutes. Formaldehyde crosslinking was quenched with 2.5 M glycine for 5 minutes. Cells were collected in PBS and incubated in nuclear lysis buffer on ice for 10 minutes (50 mM Tris pH 8.0, 10 mM EDTA, 1% SDS). Cells were sonicated using a Diagenode Bioruptor for 15 cycles in alternating 30 s on/off cycles on the low setting. Difficult to sonicate cell lines, including primary mouse colonic epithelium, were first incubated in 0.25% trypsin for 3 minutes with agitation at room temperature followed by inactivation with media containing 10% FBS. Sonicated chromatin was diluted 1:4 in IP dilution buffer (20 mM Tris pH 8.0, 2 mM EDTA, 50 mM NaCl, 1% Triton X-100, 0.1% SDS) and precleared with overheard rotation at 4 degrees for two hours using Protein A+G conjugated magnetic beads (Millipore). Chromatin was incubated with primary antibody against mouse p53 (CM5, Leica) overnight followed by two hours of conjugation to Protein A+G conjugated magnetic beads. Beads were washed once with IP Wash Buffer 1 (20 mM Tris pH 8.0, 2 mM EDTA, 50 mM NaCl, 1% Triton X-100, 0.1% SDS), twice with High Salt Wash Buffer (20 mM Tris pH 8.0, 2 mM EDTA, 500 mM NaCl, 1% Triton X-100, 0.01% SDS), once with IP Wash Buffer II (10 mM Tris pH 8.0, 1 mM EDTA, 0.25 M LiCl, 1% NP-40, 1% deoxycholic acid), and twice with TE buffer. Beads were eluted with buffer containing 1% SDS and 0.1 mM NaHCO3 with agitation at 37°C. Reverse crosslinking was performed overnight at 65^>°^C with high NaCl and RNase treatment (Roche). Proteinase K (Roche) digestion was performed for 2 hours at 42^>°^C before DNA was purified using QIAQuick PCR purification kits according to the manufacturer’s instructions.

### ChIP library preparation and sequencing

Briefly, end repair of ChIP DNA was performed using Klenow DNA Polymerase, T4 DNA Polymerase, and T4 PNK (NEB). Polyadenylation was performed using Klenow Exo (NEB). NEXTFLEX barcodes were ligated using the Quick Ligation Kit (NEB). Libraries were amplified for 15 cycles using HotStart HiFi Ready Mix (KAPA). Libraries were run on a 2% low melt agar gel, and 300-400 bp fragments were excised using the DNA Gel Extraction Kit (QIAGEN). Libraries underwent quality control using Qubit and Bioanalyzer High Sensitivity assays before being sequenced on an Illumina NextSeq 2500 at the Oncological Sciences Sequencing Core Facility at Mount Sinai. p53 ChIP libraries were sequenced to a target depth of 30 M reads and input libraries of 60 M reads.

### ChIP data analysis pipeline

Sequenced reads containing barcode contamination were filtered using Cutadapt. Reads were aligned to the mm9 mouse or hg19 human reference genome using Bowtie. Duplicated reads were removed using samtools. Peak calling was performed using MACS2 and a p-value cutoff of 1e-6. Bigwig tracks were generated and visualized on the IGV browser (Broad). Motif analysis was performed using HOMER.

### Data visualization and analysis

CyTOF data was first debarcoded using Single Cell Debarcoder^46^ using post-assignment debarcode stringency filter and outlier trimming. Clean, concatenated files were then visualized using viSNE^47^, a dimensionality reduction method, which uses the Barnes-Hut acceleration of the t-SNE algorithm. viSNE was implemented using Cytobank and generated using input tag expression levels. Cell clusters were defined by the tag. Heatmaps of cell clusters were generated by taking the median untransformed or arc-sine transformed or the percentage of negative cells within clusters and using this value unscaled or Z scaled relative to other cell clusters. For statistical analysis, following Pro-Code/gRNA assignment of CyTOF data by the single-cell debarcoder^46^, demultiplexed FCS files were read using the flowCore^48^ package in R. The resulting matrix provided protein measurements on all markers for every cell, indexed by sgRNA. For subsequent analysis, sgRNAs with less than 50 cells assigned in either the control or Cas9 conditions were removed. To determine sgRNA effects in the settings with or without IFNy, methods similar to those described in Adamson *et al*^49^ were used, where for each sgRNA all proteins measured are subjected to a Kolmogorov-Smirnov test comparing the Cas9 and negative control population empirical distribution functions of log10 intensity values. Benjamini-Hochberg correction^50^ was carried out on the resulting p-values and the effect size was measured by the magnitude of the test statistic D, the change in median fluorescence intensity (MFI), and the change in the percentage of Cas9 cells having intensity values outside the 5th or 95th percentile of control cell intensities for a given marker with the same sgRNA. Comparisons with a D value greater than 0.25 and an adjusted p-value <0.01 were deemed significant and selected for further study. GraphPad Prism 8 software was used to perform all statistical analyses. For comparison of two groups, two-tailed unpaired Student’s t tests were used. One-way ANOVA was used to compare three or more groups. Values at p < 0.05 were considered to be statistically significant.

