## Supplementary Figures for "P53 is a direct regulator of the immune co-stimulatory molecule CD80"

**Figure S1**

**a**

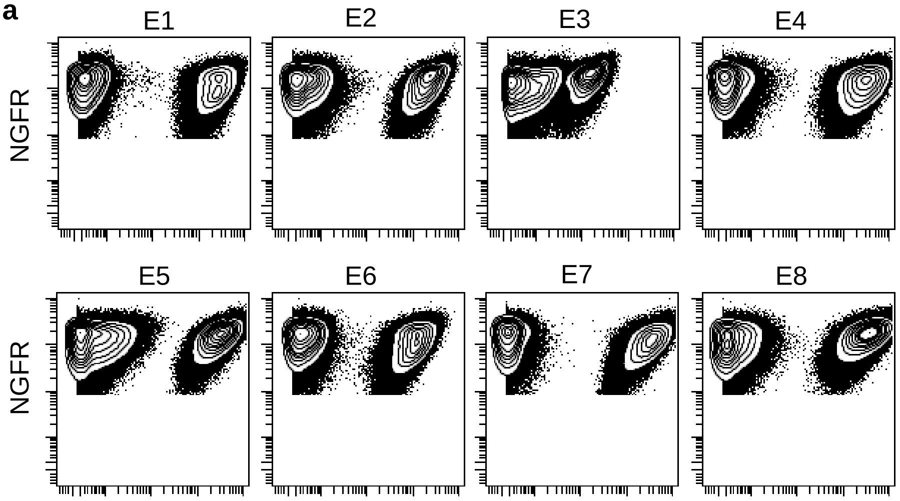


**b**

12124 Library

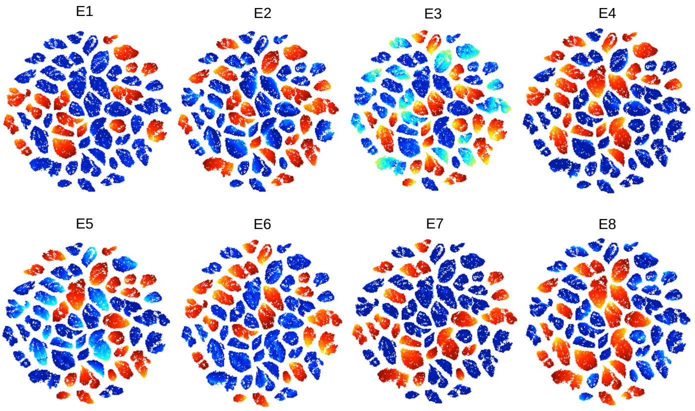


**c**

12125 Library

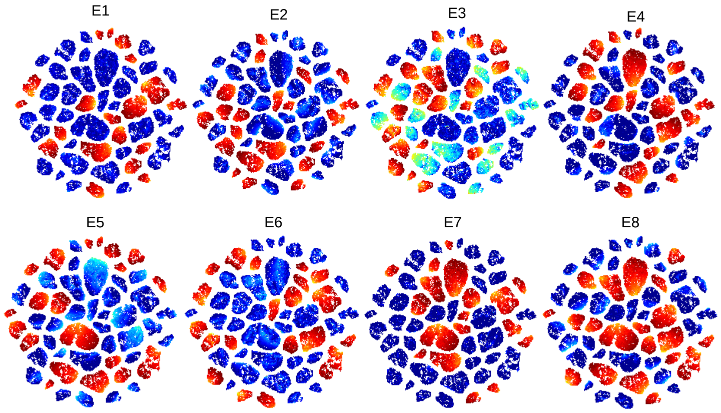

**Figure S1. Resolution and Distribution of 104 Pro-Code-Expressing Populations. Related to Figure 1.**

a. CT26 cells were transduced with 104 different Pro-Code expressing vectors, stained with metal-conjugated antibodies specific for each epitope (E1-8), and analyzed by CyTOF. X-axis represents epitope tag staining, y-axis is NGFR.

b. CT26 cells were transduced with two different 52 Pro-Code expressing vector libraries, stained with metal-conjugated antibodies specific for each epitope (E1-8), and analyzed by CyTOF. Individual viSNE plots showing expression of each of the indicated epitopes in (B) 12124 library and (C) 12125 library. Expression level is scaled from high to low (red to blue).

**Figure S2**

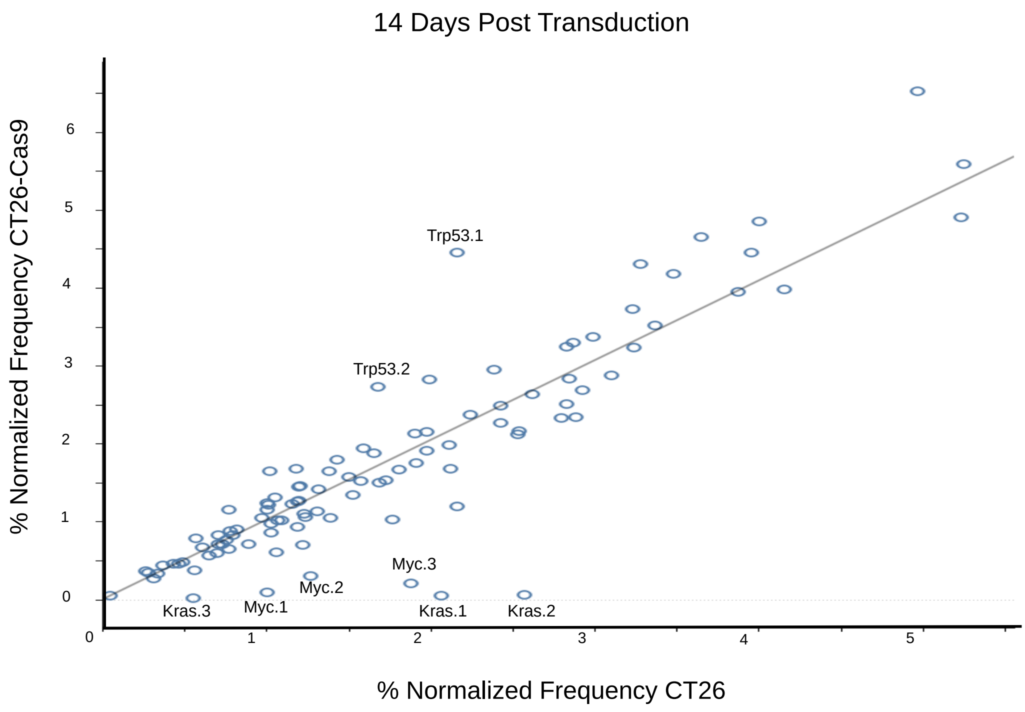
14 Days Post Transduction
% Normalized Frequency CT26-Cas9
% Normalized Frequency CT26
0
1
2
3
4
5
6
Trp53.1
Trp53.2
Myc.3
Myc.2
Myc.1
Kras.3
Kras.1
Kras.2

**Figure S2. Enrichment and De-enrichment of different Pro-Code/CRISPR engineered CT26 cells over time. Related to Figure 1.**

CT26 and CT26-Cas9 cells were transduced with 104 Pro-Code/CRISPR expressing vectors, NGFR sorted, cultured for 14 days, stained with metal-conjugated antibodies specific for each epitope (E1-8), and analyzed by CyTOF. Event counts were exported and transformed into frequency of the total population. Change in frequency >2-fold was considered significant.

### Figure S3

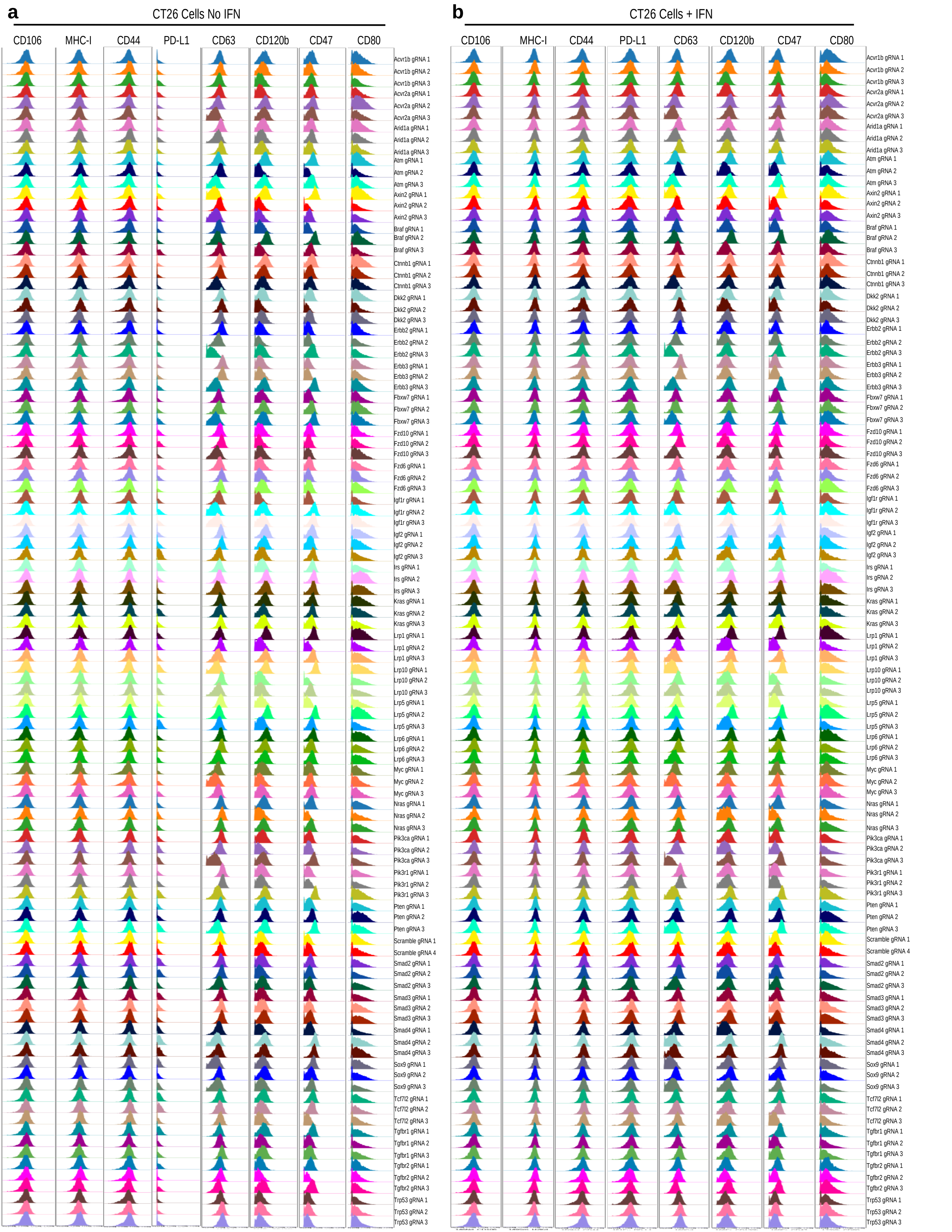
a
CT26 Cells No IFN
CD106
MHC-I
CD44
PD-L1
CD63
CD120b
CD47
CD80
Acvr1b gRNA 1
Acvr1b gRNA 2
Acvr1b gRNA 3
Acvr2a gRNA 1
Acvr2a gRNA 2
Acvr2a gRNA 3
Arid1a gRNA 1
Arid1a gRNA 2
Arid1a gRNA 3
Atm gRNA 1
Atm gRNA 2
Atm gRNA 3
Axin2 gRNA 1
Axin2 gRNA 2
Axin2 gRNA 3
Braf gRNA 1
Braf gRNA 2
Braf gRNA 3
Ctnnb1 gRNA 1
Ctnnb1 gRNA 2
Ctnnb1 gRNA 3
Dkk2 gRNA 1
Dkk2 gRNA 2
Dkk2 gRNA 3
Erbb2 gRNA 1
Erbb2 gRNA 2
Erbb2 gRNA 3
Erbb3 gRNA 1
Erbb3 gRNA 2
Erbb3 gRNA 3
Fbxw7 gRNA 1
Fbxw7 gRNA 2
Fbxw7 gRNA 3
Fzd10 gRNA 1
Fzd10 gRNA 2
Fzd10 gRNA 3
Fzd6 gRNA 1
Fzd6 gRNA 2
Fzd6 gRNA 3
Igf1r gRNA 1
Igf1r gRNA 2
Igf1r gRNA 3
Igf2 gRNA 1
Igf2 gRNA 2
Igf2 gRNA 3
Irs gRNA 1
Irs gRNA 2
Irs gRNA 3
Kras gRNA 1
Kras gRNA 2
Kras gRNA 3
Lrp1 gRNA 1
Lrp1 gRNA 2
Lrp1 gRNA 3
Lrp10 gRNA 1
Lrp10 gRNA 2
Lrp10 gRNA 3
Lrp5 gRNA 1
Lrp5 gRNA 2
Lrp5 gRNA 3
Lrp6 gRNA 1
Lrp6 gRNA 2
Lrp6 gRNA 3
Myc gRNA 1
Myc gRNA 2
Myc gRNA 3
Nras gRNA 1
Nras gRNA 2
Nras gRNA 3
Pik3ca gRNA 1
Pik3ca gRNA 2
Pik3ca gRNA 3
Pik3r1 gRNA 1
Pik3r1 gRNA 2
Pik3r1 gRNA 3
Pten gRNA 1
Pten gRNA 2
Pten gRNA 3
Scramble gRNA 1
Scramble gRNA 4
Smad2 gRNA 1
Smad2 gRNA 2
Smad2 gRNA 3
Smad3 gRNA 1
Smad3 gRNA 2
Smad3 gRNA 3
Smad4 gRNA 1
Smad4 gRNA 2
Smad4 gRNA 3
Sox9 gRNA 1
Sox9 gRNA 2
Sox9 gRNA 3
Tcf7l2 gRNA 1
Tcf7l2 gRNA 2
Tcf7l2 gRNA 3
Tgfbr1 gRNA 1
Tgfbr1 gRNA 2
Tgfbr1 gRNA 3
Tgfbr2 gRNA 1
Tgfbr2 gRNA 2
Tgfbr2 gRNA 3
Trp53 gRNA 1
Trp53 gRNA 2
Trp53 gRNA 3
b
CT26 Cells + IFN
CD106
MHC-I
CD44
PD-L1
CD63
CD120b
CD47
CD80
Acvr1b gRNA 1
Acvr1b gRNA 2
Acvr1b gRNA 3
Acvr2a gRNA 1
Acvr2a gRNA 2
Acvr2a gRNA 3
Arid1a gRNA 1
Arid1a gRNA 2
Arid1a gRNA 3
Atm gRNA 1
Atm gRNA 2
Atm gRNA 3
Axin2 gRNA 1
Axin2 gRNA 2
Axin2 gRNA 3
Braf gRNA 1
Braf gRNA 2
Braf gRNA 3
Ctnnb1 gRNA 1
Ctnnb1 gRNA 2
Ctnnb1 gRNA 3
Dkk2 gRNA 1
Dkk2 gRNA 2
Dkk2 gRNA 3
Erbb2 gRNA 1
Erbb2 gRNA 2
Erbb2 gRNA 3
Erbb3 gRNA 1
Erbb3 gRNA 2
Erbb3 gRNA 3
Fbxw7 gRNA 1
Fbxw7 gRNA 2
Fbxw7 gRNA 3
Fzd10 gRNA 1
Fzd10 gRNA 2
Fzd10 gRNA 3
Fzd6 gRNA 1
Fzd6 gRNA 2
Fzd6 gRNA 3
Igf1r gRNA 1
Igf1r gRNA 2
Igf1r gRNA 3
Igf2 gRNA 1
Igf2 gRNA 2
Igf2 gRNA 3
Irs gRNA 1
Irs gRNA 2
Irs gRNA 3
Kras gRNA 1
Kras gRNA 2
Kras gRNA 3
Lrp1 gRNA 1
Lrp1 gRNA 2
Lrp1 gRNA 3
Lrp10 gRNA 1
Lrp10 gRNA 2
Lrp10 gRNA 3
Lrp5 gRNA 1
Lrp5 gRNA 2
Lrp5 gRNA 3
Lrp6 gRNA 1
Lrp6 gRNA 2
Lrp6 gRNA 3
Myc gRNA 1
Myc gRNA 2
Myc gRNA 3
Nras gRNA 1
Nras gRNA 2
Nras gRNA 3
Pik3ca gRNA 1
Pik3ca gRNA 2
Pik3ca gRNA 3
Pik3r1 gRNA 1
Pik3r1 gRNA 2
Pik3r1 gRNA 3
Pten gRNA 1
Pten gRNA 2
Pten gRNA 3
Scramble gRNA 1
Scramble gRNA 4
Smad2 gRNA 1
Smad2 gRNA 2
Smad2 gRNA 3
Smad3 gRNA 1
Smad3 gRNA 2
Smad3 gRNA 3
Smad4 gRNA 1
Smad4 gRNA 2
Smad4 gRNA 3
Sox9 gRNA 1
Sox9 gRNA 2
Sox9 gRNA 3
Tcf7l2 gRNA 1
Tcf7l2 gRNA 2
Tcf7l2 gRNA 3
Tgfbr1 gRNA 1
Tgfbr1 gRNA 2
Tgfbr1 gRNA 3
Tgfbr2 gRNA 1
Tgfbr2 gRNA 2
Tgfbr2 gRNA 3
Trp53 gRNA 1
Trp53 gRNA 2
Trp53 gRNA 3

**Figure S3 con't**

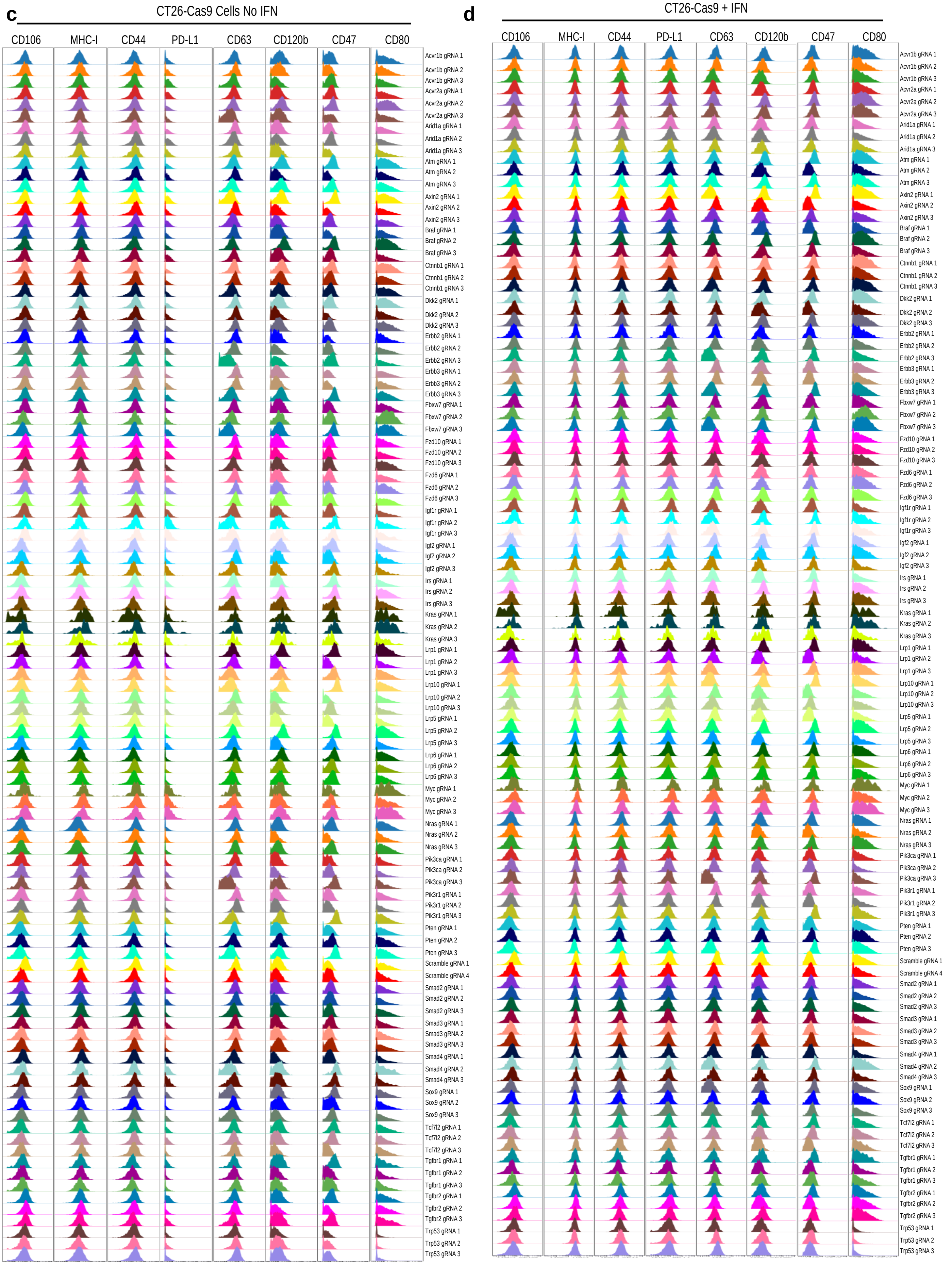

**Figure S3 con't**

**e**

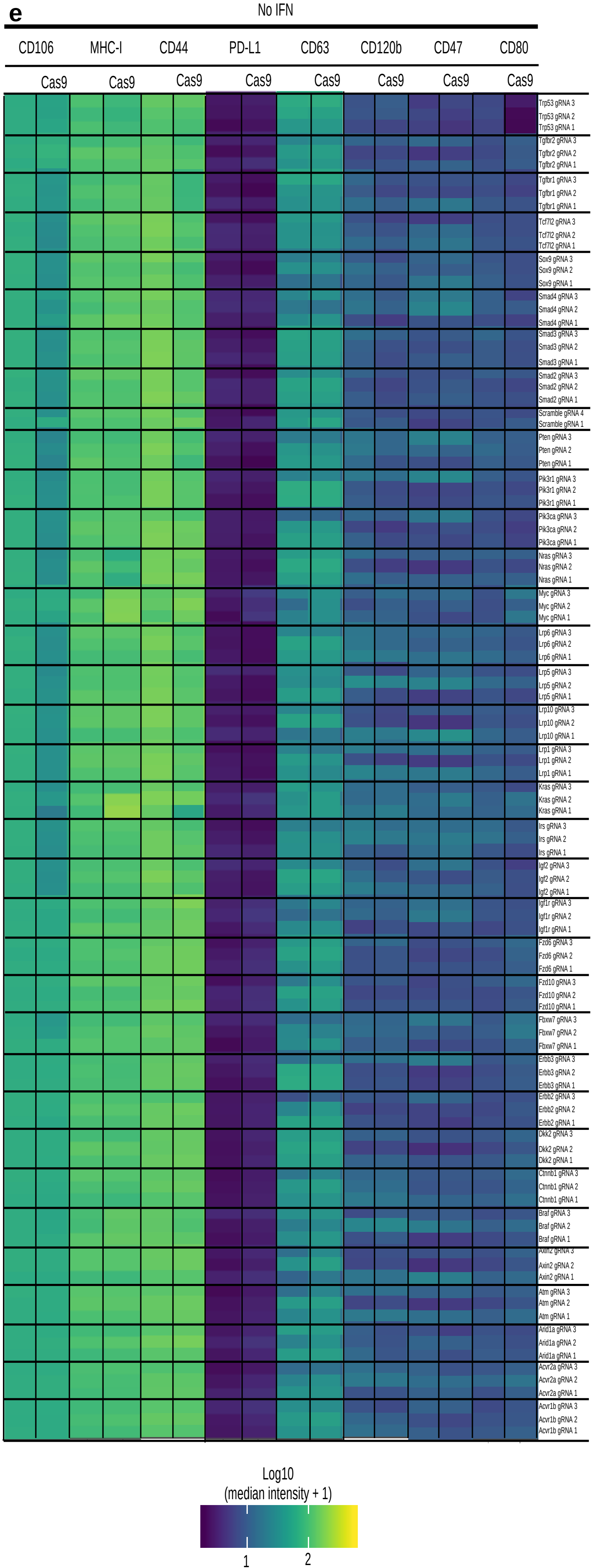


**f**

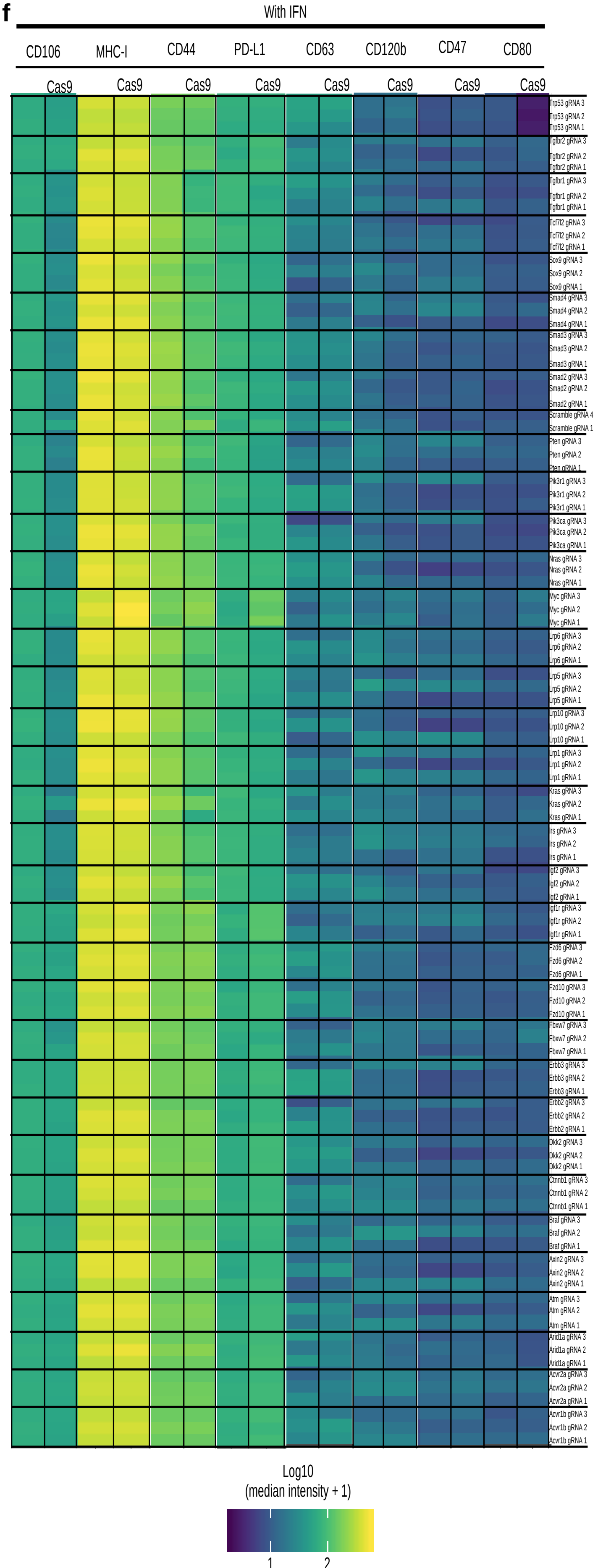

**Figure S3. Pro-Code/CRISPR screen of commonly mutated genes in colorectal cancer cells identifies multiple regulators of immunophenotype. Related to Figure 1.**

a – d. Expression of the indicated proteins on each Pro-Code/CRISPR cell population after IFNγ treatment. Shown are representative histograms for each Pro-Code population. The y axis represents cell count normalized by the protein detection channel.

e – f. Heatmap representation of the relative expression of molecules MHC-I, CD106, CD80, CD47, PD-L1, CD44, CD63, and CD120b across all Pro-Code/CRISPR populations after IFNγ treatment. All data is representative of 3 independent experiments.

**Figure S4**

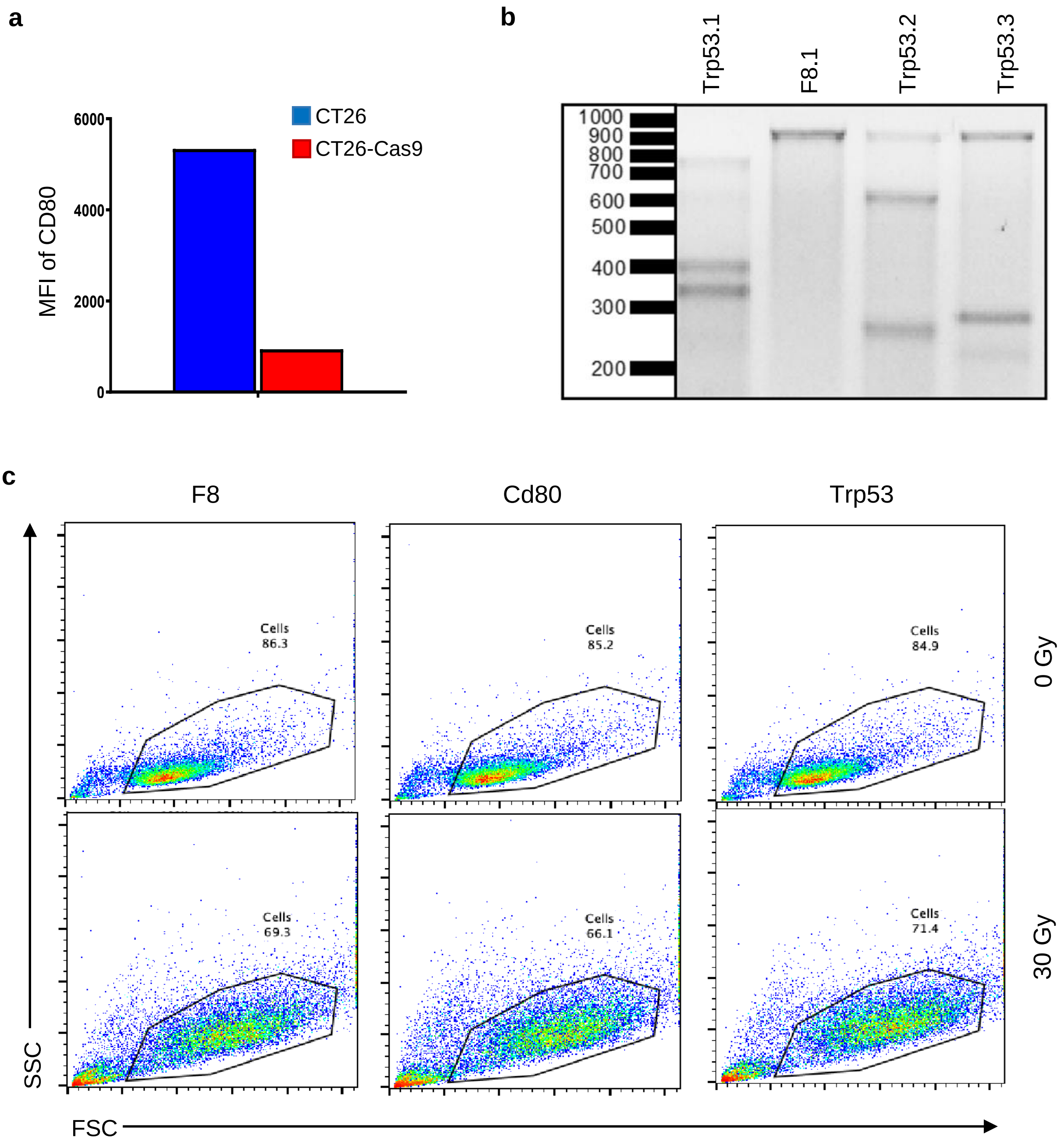
a
6000
4000
2000
0
MFI of CD80
CT26
CT26-Cas9
b
Trp53.1
F8.1
Trp53.2
Trp53.3
1000
900
800
700
600
500
400
300
200
c
F8
Cd80
Trp53
Cells
86.3
Cells
85.2
Cells
84.9
0 Gy
Cells
69.3
Cells
66.1
Cells
71.4
30 Gy
SSC
FSC

**Figure S4. TP53 positively regulates CD80. Related to Figure 2.**

a. Bar graph quantification of Figure 2a in median fluorescent intensity (MFI).

b. Surveyor mutagenesis PCR of CT26-Cas9 cells stably transduced with Trp53 or F8 gRNA vectors.

c. CT26-Cas9-GFP cells transduced individually with Trp53 gRNA.1, Cd80 gRNA.1, or F8 gRNA.1 vector were either untreated (0 Gy) or gamma irradiated (30 Gy) and analyzed by flow cytometry. Representative dot plots are shown (n = 3).

**Figure S5**

**a**

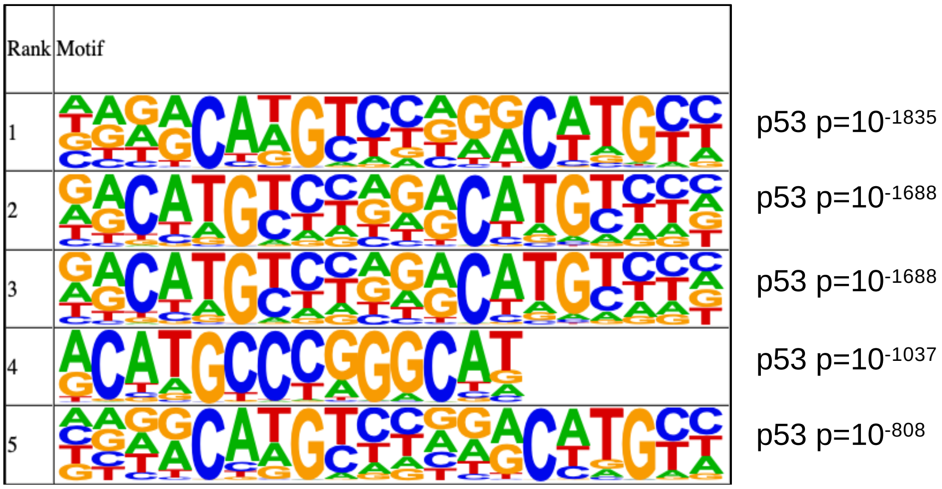
Rank
Motif
1
2
3
4
5
p53 p=10-1835
p53 p=10-1688
p53 p=10-1688
p53 p=10-1037
p53 p=10-808

**b**

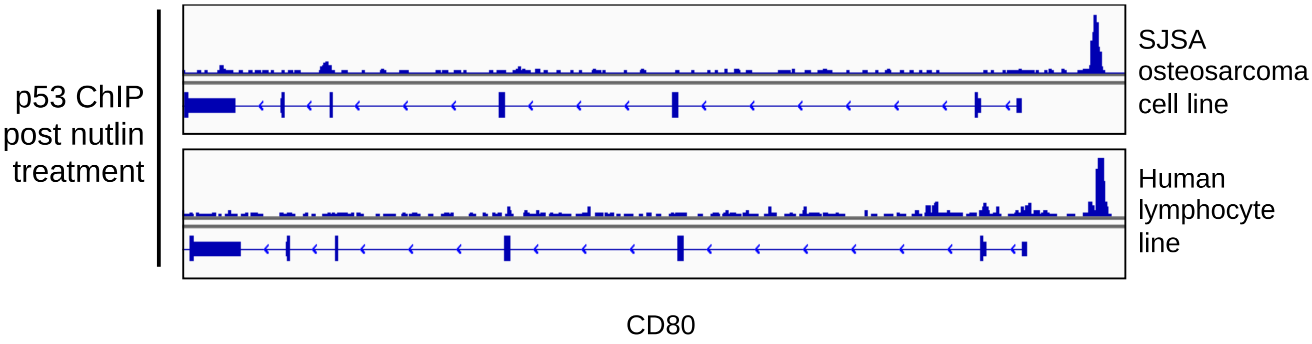
p53 ChIP post nutlin treatment
SJSA osteosarcoma cell line
Human lymphocyte line
CD80

**Figure S5. P53 is a direct regulator of CD80. Related to Figure 3.**

a. Motifs present in P53 ChIP-Seq peaks generated by MEME based on top 500 peaks called. Motif analysis was performed using HOMER.

b. P53 ChIP-Seq publicly available data post Nutlin-3 treatment was queried from Gene Expression Omnibus (GEO) and the Cistrome databases. Bigwig tracks were downloaded and visualized on the IGV browser (Broad).

**Figure S6**

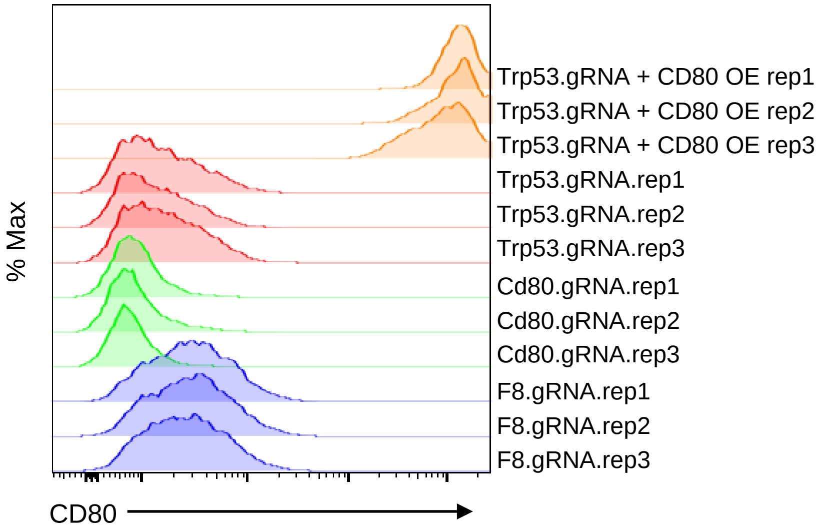
% Max
Trp53.gRNA + CD80 OE rep1
Trp53.gRNA + CD80 OE rep2
Trp53.gRNA + CD80 OE rep3
Trp53.gRNA.rep1
Trp53.gRNA.rep2
Trp53.gRNA.rep3
Cd80.gRNA.rep1
Cd80.gRNA.rep2
Cd80.gRNA.rep3
F8.gRNA.rep1
F8.gRNA.rep2
F8.gRNA.rep3
CD80

**Figure S6. Overexpression of Cd80 in p53-KO cells. Related to Figure 4.**

Representative flow cytometry histograms for CD80 in indicated CT26 modified cell lines.

#### Figure S7

**a**

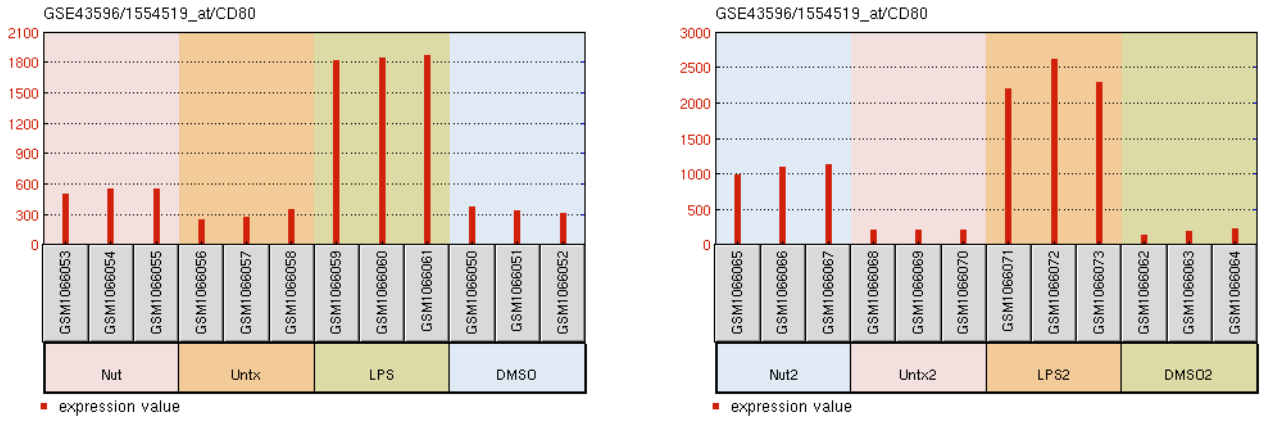


**b**

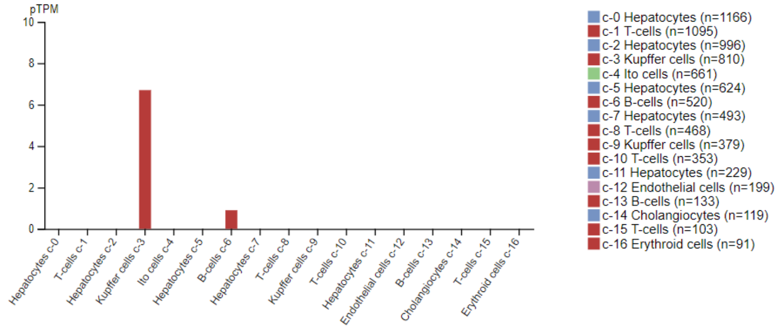


**c**

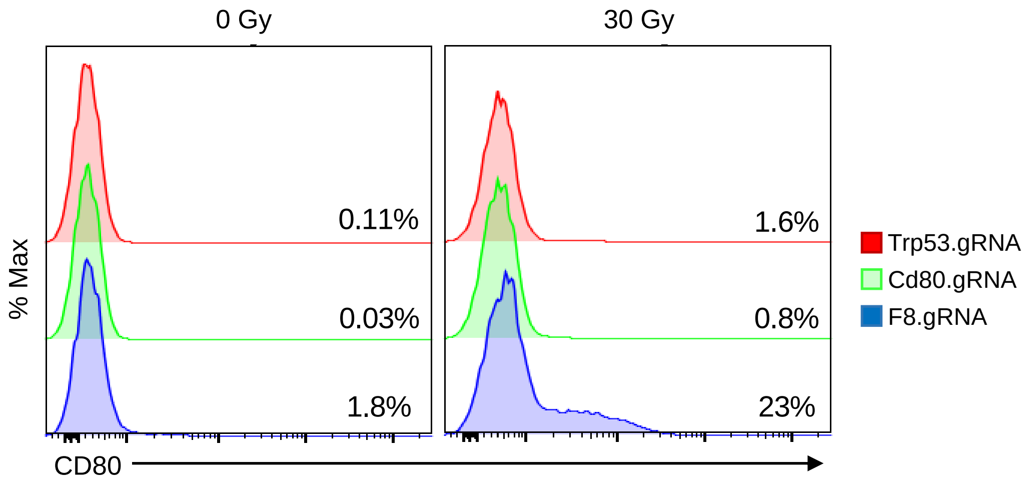

**Figure S7. Regulation of CD80 by P53 in additional cell types. Related to Figure 5.**

a. Microarray gene expression data for CD80 in human ex-vivo differentiated macrophages from two healthy donors untreated or treated with DMSO, 100 ng/ml LPS, or 10 µM Nutlin-3 for 2 hours. NCBI GEO dataset GSE43596 from Lowe JM et al. Cancer Res 2014 was analyzed with GEO2R.

b. Bar graph comparison of CD80 expression levels in liver cell types. The scRNA-sequencing data from MacParland SA et al. (2018) is visualized by the Human Protein Atlas. The read counts were normalized to transcripts per million protein coding genes (pTPM).

c. Representative flow cytometry histograms for CD80 in indicated JS1 hepatic stellate cells modified with CRISPR-CAS9 to knockout Trp53, Cd80, or F8 control and treated with 0 or 30 Gy of irradiation.
